# Rolling Circle RNA Synthesis Catalysed by RNA

**DOI:** 10.1101/2021.11.30.470609

**Authors:** Emil Laust Kristoffersen, Matthew Burman, Agnes Noy, Philipp Holliger

## Abstract

RNA-catalysed RNA replication is widely considered a key step in the emergence of life’s first genetic system. However, RNA replication can be impeded by the extraordinary stability of duplex RNA products, which must be dissociated for re-initiation of the next replication cycle. Here we have explored rolling circle synthesis (RCS) as a potential solution to this strand separation problem. RCS on small circular RNAs - as indicated by molecular dynamics simulations - induces a progressive build-up of conformational strain with destabilisation of nascent strand 5’ and 3’ ends. At the same time, we observe sustained RCS by a triplet polymerase ribozyme on small circular RNAs over multiple orbits with strand displacement yielding concatemeric RNA products. Furthermore, we show RCS of a circular Hammerhead ribozyme capable of self-cleavage and re-circularisation. Thus, all steps of a viroid-like RNA replication pathway can be catalysed by RNA alone. Our results have implications for the emergence of RNA replication and for understanding the potential of RNA to support complex genetic processes.

## Introduction

The versatility of RNA functions underpins hypotheses regarding the origin and early evolution of life. Such hypotheses of an “RNA world” – a primordial biology centred on RNA as the main biomolecule - are in accord with the essential role of RNA catalysis in present day biology (Cech, 2000; Goldman and Kacar, 2021; Nissen et al., 2000; Wilkinson et al., 2020) and the discovery of multiple prebiotic synthetic pathways to several of the RNA (and DNA) nucleotides (Becker et al., 2019; Kim et al., 2020; Patel et al., 2015; Powner et al., 2009; Xu et al., 2020). In addition, progress in both non-enzymatic (Deguzman et al., 2014; Hassenkam et al., 2020; Prywes et al., 2016; Rajamani et al., 2008; Wachowius and Holliger, 2019; Zhang et al., 2020; Zhou et al., 2020) and RNA-catalysed polymerization of RNA and some of its analogues (Attwater et al., 2018, 2013; Cojocaru and Unrau, 2021; Ekland and Bartel, 1996; Horning and Joyce, 2016; Johnston et al., 2001; Mutschler et al., 2018; Shechner et al., 2009; Tagami et al., 2017; Tjhung et al., 2020) is beginning to map out a plausible path to RNA self-replication; a cornerstone of the RNA world hypothesis.

RNA *in vitro* evolution and engineering have led to the discovery of RNA polymerase ribozymes (RPRs) able perform templated RNA synthesis of up to ∼200 nucleotides (nt) (Attwater et al., 2018), synthesizing active ribozymes including the catalytic class I ligase core (Horning and Joyce, 2016; Tjhung et al., 2020) at the heart of the most efficient RPRs, as well as initiate processive RNA synthesis using a mechanism with analogies to sigma-dependent transcription initiation (Cojocaru and Unrau, 2021). A RPR capable of utilizing trinucleotide triphosphates (triplets) as substrates (a triplet polymerase ribozyme (TPR)) has been shown to display a much enhanced capacity to copy highly structured RNA templates including segments of its own sequence (Attwater et al., 2018).

Nevertheless, there remain a number of fundamental obstacles to be overcome before an autonomous self-replication system can be established. A central problem among these is the so called “strand inhibition problem”, a form of product inhibition due to the accumulation of highly stable dead-end RNA duplexes, which cannot be dissociated (efficiently) under replication conditions (Le Vay and Mutschler, 2019). The strand inhibition problem has been overcome by (PCR-like) thermocycling (or thermophoresis) (Horning and Joyce, 2016; Salditt et al., 2020) but this approach may be limited to short RNA oligomers (even in the presence of high concentrations of denaturing agents) as the melting temperatures of longer RNA duplexes approach or even exceed the boiling point of water (Freier et al., 1986; Szostak, 2012).

While RNA duplexes occur by necessity as intermediates of RNA replication, the extent of the strand inhibition problem can be modulated by genome topology. Circular rather than linear genomes are widespread in biology including eukaryotes, prokaryotes and viruses (Møller et al., 2018; Moss et al., 2020; Shulman and Davidson, 2017). Circular RNAs (circRNAs) are found as products of RNA splicing (Kristensen et al., 2019) and RNA-based self-circularization is known in multiple ribozymes (Hieronymus and Müller, 2019; Lasda and Parker, 2014; Petkovic and Müller, 2015). Templated RNA synthesis on circular templates (Rolling Circle Synthesis (RCS)) is also widespread and found in the replication of the RNA genomes in some viruses and in viroids. Indeed, viroid RNA replication has been proposed to resemble an ancient mechanisms for replication (Diener, 1989; Flores et al., 2014). RCS has potentially unique properties with regards to the strand inhibition problem where RNA duplex melting in principle can be effected by continuous toehold strand displacement driven by nucleotide hybridization and the ratchet of nascent strand extension by triphosphate hydrolysis. In an idealized RCS mechanism, such strand invasion and displacement processes are both isoenergetic and coordinated to nascent strand extension (Blanco et al., 1989; Daubendiek et al., 1995), with rotation of the single-stranded RNA (ssRNA) preventing the build-up of topological tension (Kuhn et al., 2002). Thus RCS is a potentially open-ended process leading to the synthesis of single-stranded multiple repeat products (concatemers) with an internally energized strand displacement circumventing the “strand inhibition problem” (Tupper and Higgs, 2021).

Here we have explored RCS of small circular RNA (scRNA) templates as a potential solution to the strand inhibition problem in RNA-catalysed RNA replication. We show that RCS can be catalysed by a TPR, which is able to perform continuous templated extension of circular RNA templates for multiple cycles yielding concatemeric repeat products. We also explore the mechanistic basis for RCS and strand displacement by molecular dynamics (MD) simulations of scRNA in explicit solvent. Finally, we explore the potential of a full viroid–like replication cycle catalysed by RNA by design and synthesis of a circular Hammerhead ribozyme capable of both product cleavage and self-circularization.

## Results

### RNA-catalysed primer extension using small circular RNA templates

We first set out to investigate whether templated RNA synthesis on scRNAs could be catalysed by an RNA catalyst. To extend beyond the full circle and initiate RCS requires duplex invasion and displacement of the original RNA product strand. However, most RPRs are inhibited by duplex RNA both in the form of template secondary structures and as linear duplex RNA. We therefore explored the potential of a recently described TPR (Attwater et al., 2018), which is able to utilize triphosphorylated trinucleotides (triplets (^ppp^NNN)) as substrates for polymerization. Due to increased binding of the triplets to the template (compared e.g. to the canonical mononucleotide triphosphates (NTPs)), triplets are able to invade and cooperatively “open up” template secondary structures for replication (Attwater et al., 2018). We hypothesized that this ability might also promote the continuous invasion of the opposing strand and facilitate the RCS mechanism (Figure 1A). Similar to what was described previously, RNA synthesis by the TPR best in the eutectic phase of water ice, due to beneficial reaction conditions for ribozyme catalysis such as reduced RNA hydrolysis and high ionic and RNA substrate concentrations (Attwater et al., 2010). This was also the case on scRNA templates.

**Figure 1.**
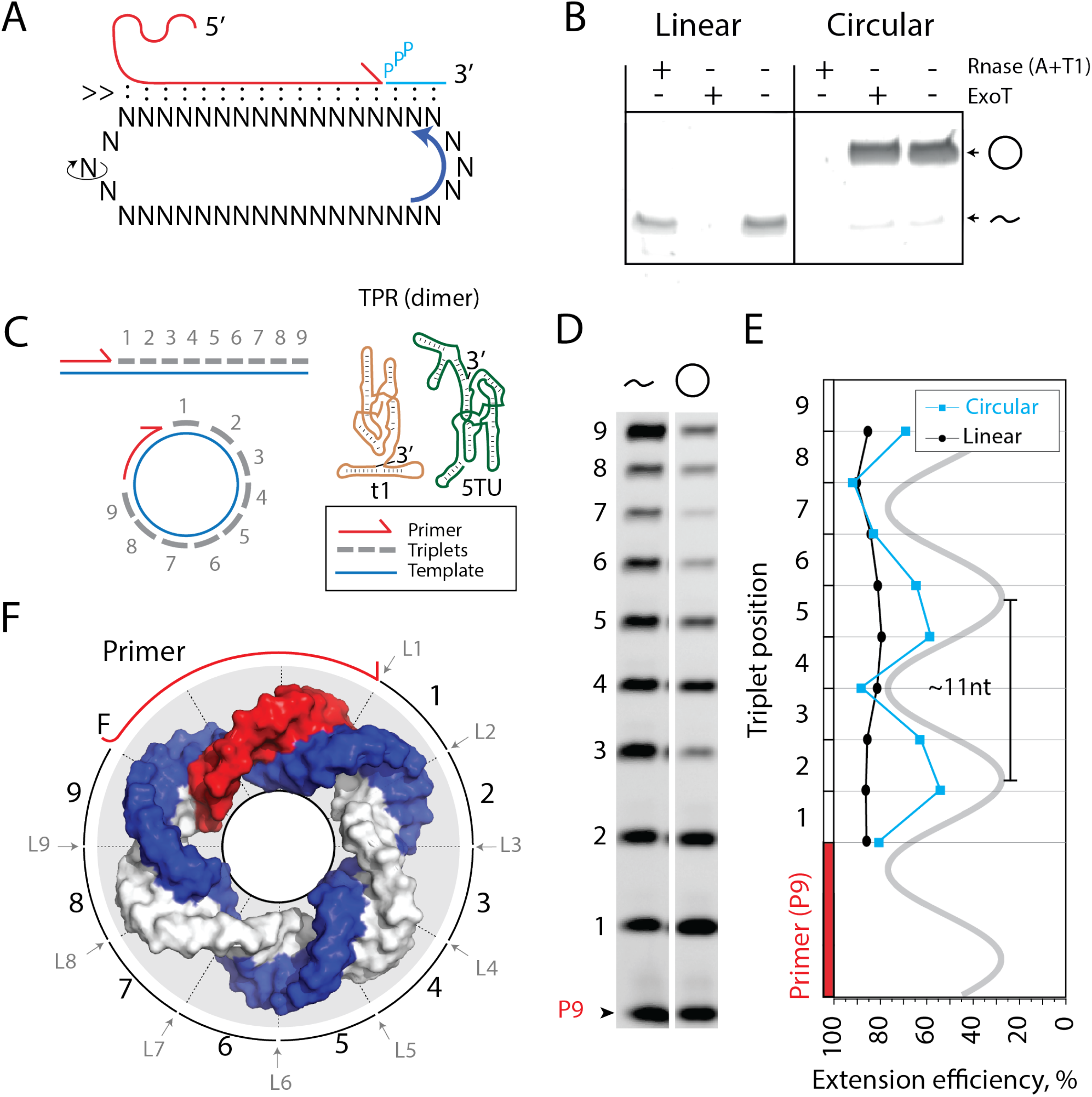
Primer extension on circular RNA templates. A) Schematic illustration of the RCS mechanism. Red product RNA strand is extended by a triplet in the 3’ end while the 5’ end dissociates by three base pairs keeping the total hybridization energy constant. Topological relaxation is allowed by rotation in the single stranded part of the circular template illustrated by swivelling arrow. B) Linear or circularized RNA is treated with or without endo- or exonucleases (RNase A/T1 mix or Exonuclease T, respectively). C) Primer extension scheme by the TPR on a linear or circular RNA template. D) PAGE gel of TPR primer extension, P9 is the unextended (fluorophore labelled) primer, bands 1-9 denote extension of P9 by 1-9 triplets ,9 extensions being full length. E) Extension efficiency of formation of band 1-9 in D) (see Materials and Methods) is plotted against triplet position. F) Schematic model of scRNA illustrating the different accessibility of in- or outside facing ligation junctions showing the scRNA template (blue), P9 primer (red) and the product strand (light grey). Original gel images and numerical values are supplied in Figure 1-source data 1.

We prepared scRNA templates (34-58 nt in length) by *in vitro* transcription and ligation and confirmed circularity by resistance to exonuclease degradation in contrast to the linear, non-ligated counterparts (Figure 1B, Figure 1 - Figure supplement 1, see sequences for all oligonucleotides in Supplementary file 1). On these, we first investigated primer extensions using just a single triplet (^ppp^GAA) as this provides an even banding pattern of incorporation facilitating analysis allowing primer extension efficiencies of linear and circular templates to be more readily compared (Figure 1C). Primer extension experiments using a purified 36 nt scRNA as template resulted in full-length extension around the circle (Figure 1D), but with reduced efficiency compared to a linear RNA template. Furthermore, we observed a periodic pattern of extension efficiency for the triplet junctions in agreement with the helical pitch of double-stranded RNA (dsRNA) (11.3 base pairs (bp)/turn (Bhattacharyya et al., 1990)) (Figure 1E). Presumably, triplet junctions located on the inside of the scRNA ring are less well accessible and therefore less efficiently ligated than in linear RNA, which is freely accessible from all sides (Figure 1F). In turn, this leads to the observed periodicity and reduced synthetic efficiency on scRNAs.

Despite the reduced extension efficiency in scRNA, we obtained full circle extension products for multiple templates (34 to 58 nt in size, Figure 2A, B) with a clear trend towards increasing mean extension efficiency for circular templates with increasing size predicting parity with the linear template at around 120 nt (Figure 2C). Note, in these experiments extension beyond full circle was not intended or possible (lane 1 to 6 in Figure 2B) as the specific triplet substrates needed for displacing the primer were not present in the reaction.

**Figure 2.**
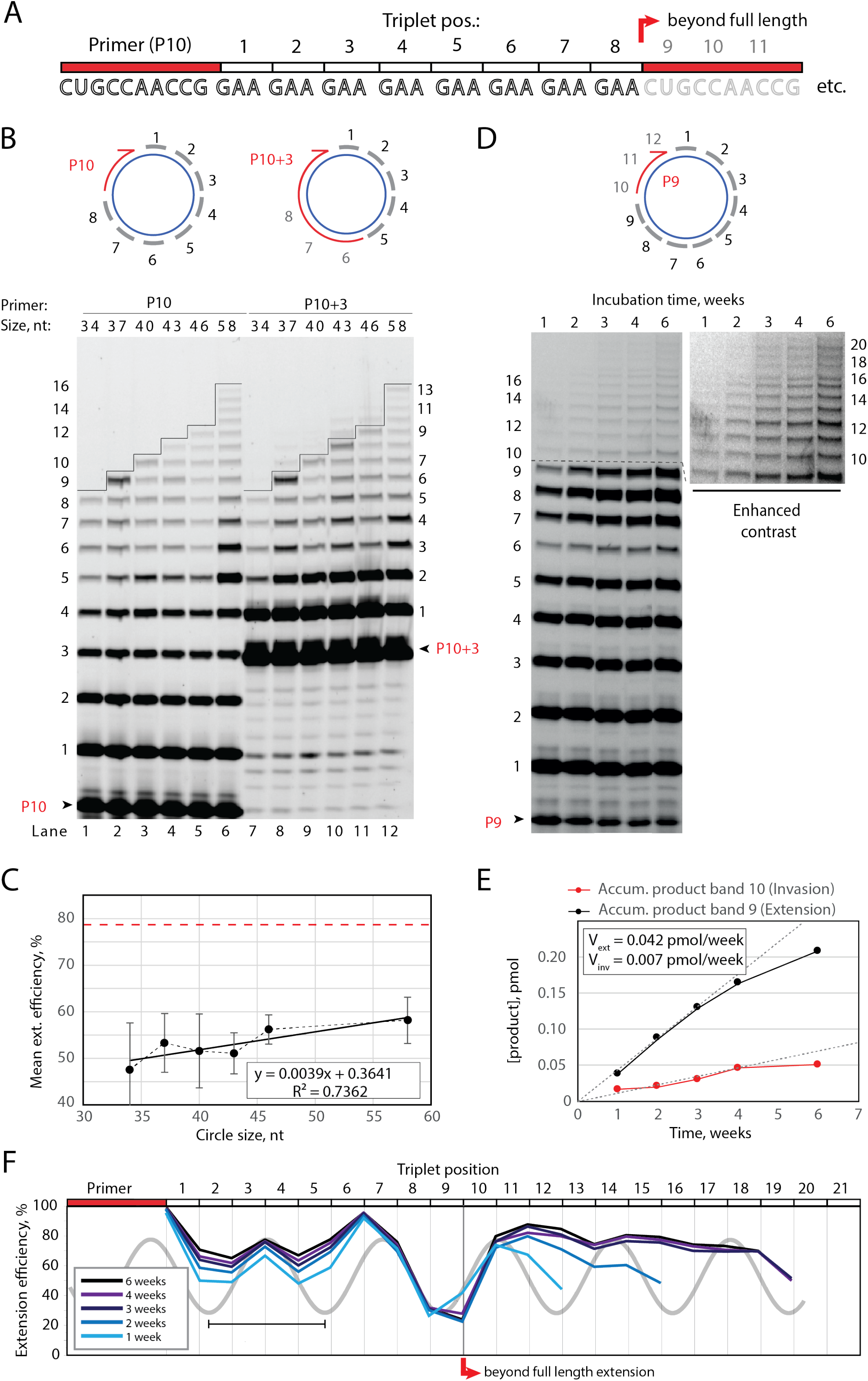
Full-length and beyond full-length RNA-catalysed RNA synthesis on circular RNA templates. A) Product strand of primer extension experiments with primer P10 (red) and 8 triplet scRNA template strand. Potential beyond full-length synthesis is shown as opaque. B) Various scRNA template sizes allow full-length primer extension as indicated (with 8 triplet sites) (blue), GAA triplets (black) and primers P10 (FAM-CUGCCAACCG) or P10+3 (FAM-CUGCCAACCG-GAA-GAA-GAA) (red). PAGE of primer extensions (under standard conditions) with full-length synthesis for different scRNA templates marked by a black line. C) Mean extension efficiency plotted as a function of circle size calculated from extension experiments including B) (Error-bars indicate standard deviation, n=5), with mean extension efficiency for a linear RNA template (red dashed line). D) scRNA template 36 nt 12xUUC-repeat and primer P9 and PAGE of time-course of primer extension (optimized conditions). Thin black line (after band 9) marks full circle synthesis. Bands 10+ (see enhanced contrast gel) indicate beyond full-length synthesis (invasion). E, F) Mean extension efficiency (from gel in D) plotted against time (E ) or triplet position (F) showing the respective amounts of product at full (black) and beyond full circle (red) synthesis as well as the efficiency drop at full length, which recovers once beyond full-length synthesis is initiated. Vext and Vinv denotes the calculated velocity of formation of band 9 and 10, respectively. Original gel images and numerical values are supplied in Figure 2-source data 1.

Having established full-length synthesis on scRNA templates, we next tested if primer extension could proceed beyond full circle requiring duplex invasion and displacement of the primer / product strand. We first tested this using primer P10+3, comprising a 5’ extension of three GAA repeats, thus covering the last three UUC triplet binding sites on the circular templates (Figure 2B top right). We observed a extension of up to three bands above the full circle mark (Figure 2B lane 7-12), indicating displacement of the primer 5’-end upon incorporation of three additional pppGAA triplets. This showed that “beyond full circle” synthesis including strand displacement is possible on scRNA templates boding well for the implementation of full RCS. To that effect, we next optimized buffer and extension conditions for more efficient extension above the full circle mark (Figure 2 - Figure supplement 1). Interestingly, greater dilution of reaction mixtures prior to freezing resulted in more efficient stand displacement. Greater dilution does not alter the final solute concentrations within the eutectic phase (Attwater et al., 2010) but increases the eutectic phase / ice interface area. This suggest that strand invasion may be aided by surface effects, as previously suggested for RNA refolding (Mutschler et al., 2015). Under these optimized buffer and extension conditions, we observed progressive accumulation of longer and longer RCS products, over prolonged reaction times (up to 6 weeks) (Figure 2D) with reaction speed decreasing after ca. 4 weeks incubation, indicating continued RCS over extended periods of time (Figure 2E, F).

### Molecular dynamics simulations of 36 nt scRNA

To better understand the structural and topological constraints of RCS on scRNAs, we performed atomistic MD simulations over 400 ns of the different RCS stages, comprising the starting scRNA template as circular single-stranded RNA (ssRNA) and scRNA with a progressively extended double-stranded RNA (dsRNA) parts (Figure 3). For simplicity, a 36 nt circular RNA sequence of (UUC)_12_ was chosen as a template strand (similar to the scRNA template in Figure 1, 2D) for direct comparison with the experimental system. The complementary strand comprising GAA triplets starting from 9 bp dsRNA (corresponding to binding of primer P_9_) was extended (in triplet increments) from 18, 21, 24, 27 till 30 bp of dsRNA (corresponding to extension products in bands 3, 4, 5, 6 and 7 in the gel in Figure 1D), using the most representative structure of the previous simulation as a starting point for the next one.

**Figure 3.**
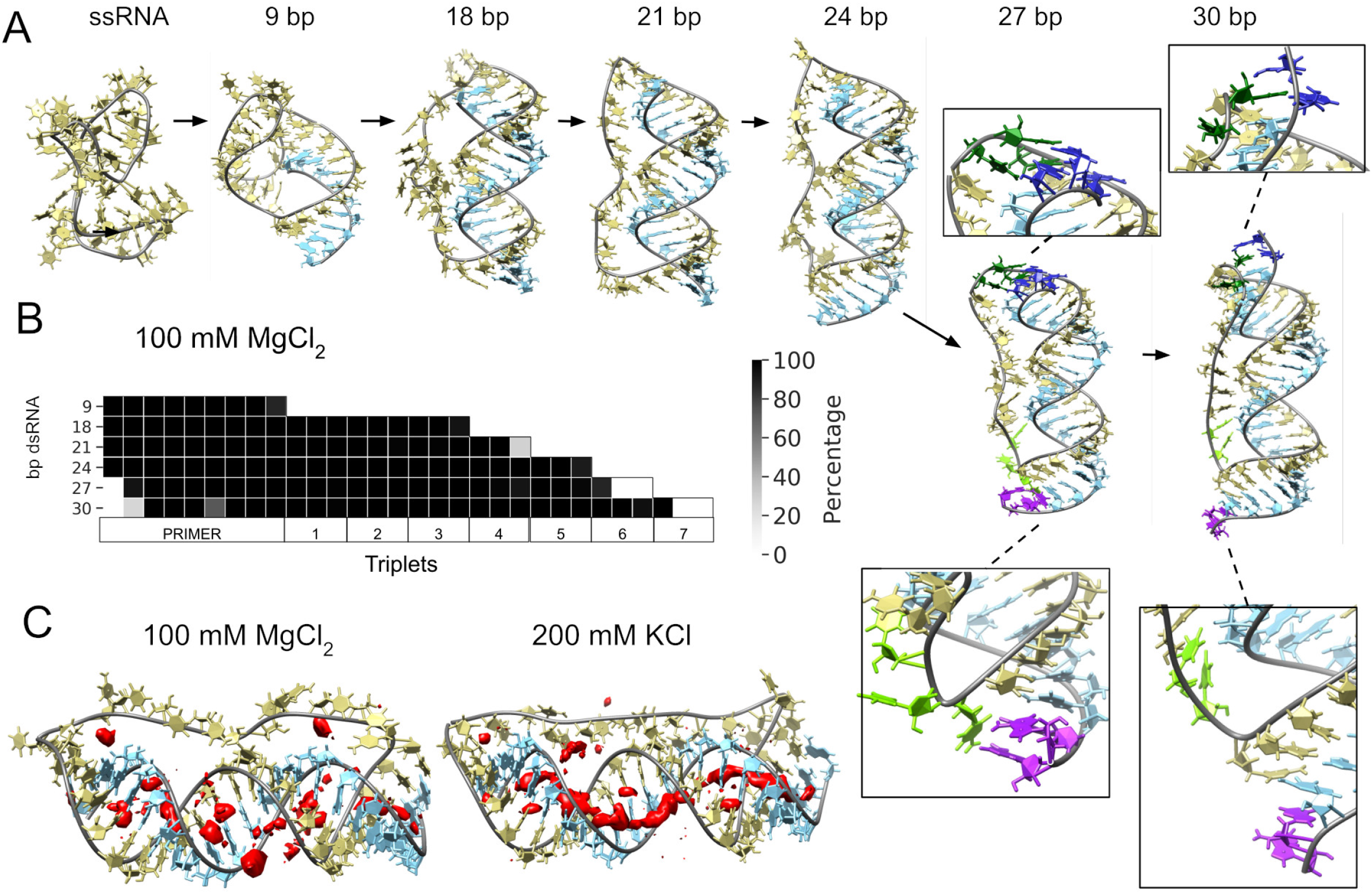
Molecular dynamics simulation of circular RNA. A) Main conformations (and zoom-in to relevant regions (squares)) observed from simulations in 100 mM MgCl2 on scRNA exploring consecutive states of primer extension, from 9 to 30 bp dsRNA with pyrimidine (template) strand (UUC)12 (khaki), purine (product) strand (GAA) (light blue), 5’ end and unpaired bases (dark blue) and 3’ end unpaired bases (purple) and matching melted bases from the template strand (dark green (5’ end) / light green (3’ end)). B) Percentage of frames from the last 100 ns of the simulations presenting canonical hydrogen bond pairing for each bp. C) Counterion-density maps (in red) around RNA molecules that show an occupancy ∼10 times or greater than the bulk concentration.

The simulation trajectories revealed the high energy barrier of dsRNA for bending and accommodating a circular shape (Figure 3A). Instead, we observe that, as dsRNA is elongated, the remaining ssRNA segment of the scRNA becomes increasingly extended. As the dsRNA part reaches 27 bp (corresponding to band 6 in Figure 1C), the ssRNA segment was fully extended and torsional strain was relieved by dissociation (“peeling off”) of the dsRNA 5’ and 3’ ends rather than by bending or the introduction of kinks into the dsRNA segment (Figure 3B). Subsequently, multiple peeling off and rebinding events were observed during the trajectories indicating that the dynamics of this process are fast (Supplementary Movie 1 and 2).

In the experimental data, we observed a larger than expected inhibitory effect for insertion of the final triplets (extension to 33 and 36 nt of dsRNA, bands 8 and 9 in Figure 2D into the corresponding scRNA template). This may reflect the onset of the 3’ and 5’ end destabilization observed in simulations (Figure 3), which would likely attenuate primer extension by the ribozyme. Our data shows a further slowdown in triplet incorporation when RCS is extended beyond full circle. We hypothesize that this might be caused by rebinding of the displaced strand on the template and interference with ribozyme extension. According to our simulations, the displaced strand would be >9 nt and, thus, long enough to reach the template strand and hybridize to the complementary repetitive sequence.

As a control for the observed dsRNA end destabilization mechanism, we also ran a simulation of a linear RNA molecule containing four triplets and a nick between two of them, but observed neither base opening nor dissociation at any strand end (Figure 3 - Figure supplement 1). Groove dimensions and local helical parameters (roll, twist and slide) for the RCS simulations on circular RNA did not show any major adjustment compared with the linear RNA control (Figure 3 - Figure supplement 2). We observed an oscillation of high / low values of bending along the molecule in phase with RNA-turn periodicity in an attempt to create an overall curvature (Velasco-Berrelleza et al., 2020), although with moderate success (∼60° on an arc length of 30 bp of dsRNA) and no formation of kinks or other disruption of the canonical A-form typical of the RNA duplex (Figure 3 - Figure supplement 2).

To mirror the experimental eutectic phase conditions, simulations were run at relatively high Mg^2+^ concentrations (100 mM) and compared with the presence of monovalent ions like K^+^ (200 mM) and high concentration of Mg^2+^ (500 mM), but simulations did not show any major differences in terms of melting or dsRNA bending (Figure 3 - Figure supplement 1, 2). However, Mg^2+^ compared to K^+^ makes more stable interactions with different parts of the RNA and, consequently, may increase the probability of distorted conformations facilitating the exposition of nucleobases at the 5’ and 3’ ends. On the contrary, K^+^ counter ions are mainly positioned along the major and minor groove, allowing the bases to orient towards the inside of the dsRNA helix for base-pairing interactions (Figure 3C and Figure 3 - Figure supplement 3). The role of Mg^2+^ in the stabilization of complex RNA folding has been observed repeatedly in several structures (Sponer et al., 2018), like the ribosome (Klein et al., 2004) and the Hepatitis delta virus ribozyme (Nakano et al., 2001). Increasing MgCl_2_ concentration to 500 mM does not seem to bring extra benefit, as the system appears to be saturated already at 100 mM Mg^2+^ (Figure 3 - Figure supplement 3, 4).

In summary, our simulations support the notion that a circular RNA template (in the presence of Mg^2+^ ions) leads to increased dynamics of nucleobase exposure, RNA duplex destabilization and 5’ and 3’ end melting, which may facilitate strand displacement during RNA replication. The simulations clearly show the implausibility of a small circular fully dsRNA molecule (as schematically illustrated in Figure 1E) due to the prohibitive energetic cost of bending of the dsRNA. Instead, the system appears to relieve internal strain by extending the ssRNA segment of the circle (partially shielding the dsRNA segment) and peeling of both dsRNA 5’ and 3’ ends (Figure 3), consistent with the helical period of triplet extension observed (Figure 1, 2) (with ligation junctions facing into the ssRNA centre being less accessible) and the observed reduction in RCS efficiency. Dynamic destabilization of dsRNA 5’ ends may aid continuous extension of the 3’ end (RCS) and would be predicted to manifest itself in RNA circles up to 200 bp as suggested by RNA persistence length (Abels et al., 2005).

### Templated rolling circle RNA synthesis

Having validated RNA synthesis on scRNA templates (Figure 1, 2) we next sought to establish RCS beyond a single “orbit” involving displacement of the primer and nascent strand. To this end, we designed barcoded templates that would allow us to distinguish TPR-made RNA products arising from non-templated terminal transferase (TT) activity from those from templated RCS by sequencing. The barcoded small RNA templates (termed A-D) were prepared either as circular or linear RNAs comprising different internal triplet “barcodes” (at position 3, 6 and 9) of variable GC-content for individual identification (Figure 4A and Figure 4 - Figure supplement 1). On these, we performed one-pot primer extension experiments, in which all four templates (either A-D linear or A-D circular) were mixed in equal proportions. After gel electrophoresis, the area above full-length extension products were excised, RNA recovered, and sequenced (Figure 4 - Figure supplement 1).

**Figure 4.**
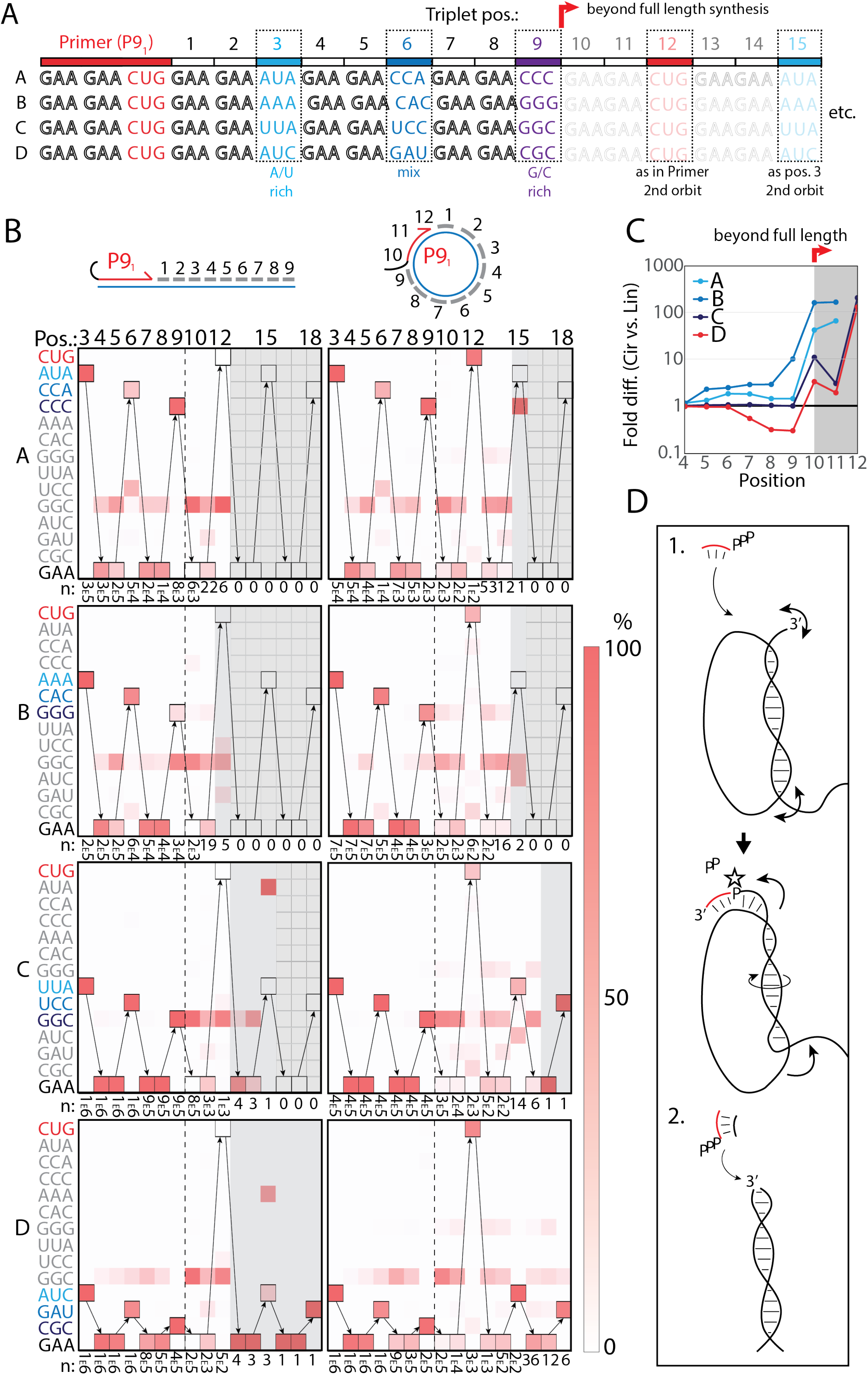
RNA-catalysed RNA synthesis beyond full length for circular templates. A) Product strands of primer extension experiments with linear and scRNA templates A-D with primer P91. Opaque sequence illustrate potential beyond full-length synthesis on scRNA. Barcode triplets at positions 3 (A/U rich) (cyan), 6 (mix) (blue), 9 (G/C rich) (purple) allow identification of product RNAs. Barcode triplet at position 15 (scRNA) is the same as that of position 3 but after one orbit on the circular template. B) Fidelity heat-map of the sequences derived from the one-pot experiments with linear (left) or circular (right) templates. Red colour indicates high prevalence of a given triplet (vertical axis) at the position noted (3-18). n: denotes the number of recovered sequence reads at each position. Transparent grey boxes cover positions with n≤5. C) Plot shows ratio (fold difference) of the probability of a product of reaching positions 4-12 on circular compared to linear templates. D) Model illustrating (1.) beyond full-length extension on a circular template (templated RCS) and (2.) on a linear template (non-templated). Full analysis of the data in Figure 4B is supplied in Figure 4-source data 1.

Analysis of the sequencing products from the one-pot reaction showed template-dependent high-fidelity RNA synthesis up to full length (position 9) for all templates (linear and circular) (Figure 4B). Further, all templates gave longer than full-length products indicative of continued RNA synthesis by the TPR after full length (positions >9). However, the fidelity dropped after full length indicative of significant non-templated terminal transferase-like (TT) activity in this regime (Figure 4B). For example, the average fidelity for insertion of the expected triplet (^ppp^GAA) for position 10 (full length +1) for circular templates was 10.9% whereas for position 9 (full length) it was 89.9%. For linear templates, the fidelity for full length +1 dropped to 0.7% compared to full length 78.8%. Note, that fidelity at full length +1 dropped much more for linear than for circular templates. For this reason, the probability of a product extending to longer than full length (positions 10-12) with the correct sequence was many fold higher for circular compared to linear RNA templates (Figure 4C). A few events of blunt-end ligation with other template / product strands (see e.g. position 15 for linear template C and D) (Figure 4B) were also observed for linear templates.

On all circular templates (with the exception of template B, where too few reads were obtained) extension beyond full length (while containing a significant TT component) continued to insert the barcode triplets correctly, indicating continuous RCS at least up to position 18 (63 nt, more than 1.5 “orbits” on the scRNA template).

Control experiments, with individually incubated templates (in contrast to the one-pot experiments) mixed after gel purification, showed essentially identical results (Figure 4 - Figure supplement 2A). Interestingly, non-diluted samples had a decreased fidelity at position 10 (the point of strand invasion) compared to diluted samples (Figure 4 - Figure supplement 2B) suggesting that dilution appears to aid not only extension efficiency (Figure 2 - Figure supplement 1D), but also strand invasion fidelity and continued templated synthesis. In summary, these results are consistent with RNA-catalysed RCS on scRNA templates beyond the full circle.

### Multiple repeat rolling circle products

Next we sought to test if RCS efficiency could be increased by double priming on the circular template, an approach known as branched RCS (Berr and Schubert, 2006). Indeed, we observed a higher degree of RCS with a 36 nt scRNA template (8211) having two identical primer sites leading to two different products being formed (product I or II) (Figure 5A, B). In order to test the primer site functionality individually we used different triplet combinations (Figure 5B). When only the ^ppp^GAA triplet was present, primers were extended only by two triplets as expected (lane 1 in Figure 5B) (with a small amount of non-templated TT incorporation of a 3rd triplet). When ^ppp^GAA and ^ppp^AUA or ^ppp^GCG triplets were added, respectively (lane 2 or 3 in Figure 5B), products extended up to 5 triplet-incorporations with extension stopping at triplet 6 (coding for CUG) showing that both primer sites were functioning. Finally, when all triplets (^ppp^GAA, ^ppp^AUA, ^ppp^GCG, ^ppp^CUG) were present, extension continued to beyond full circle (positions ≥10) (Figure 5B) and bands corresponding to extension products exceeding two whole orbits (> triplet 21 (63nt)) of replication were observed (Figure 5C).

**Figure 5.**
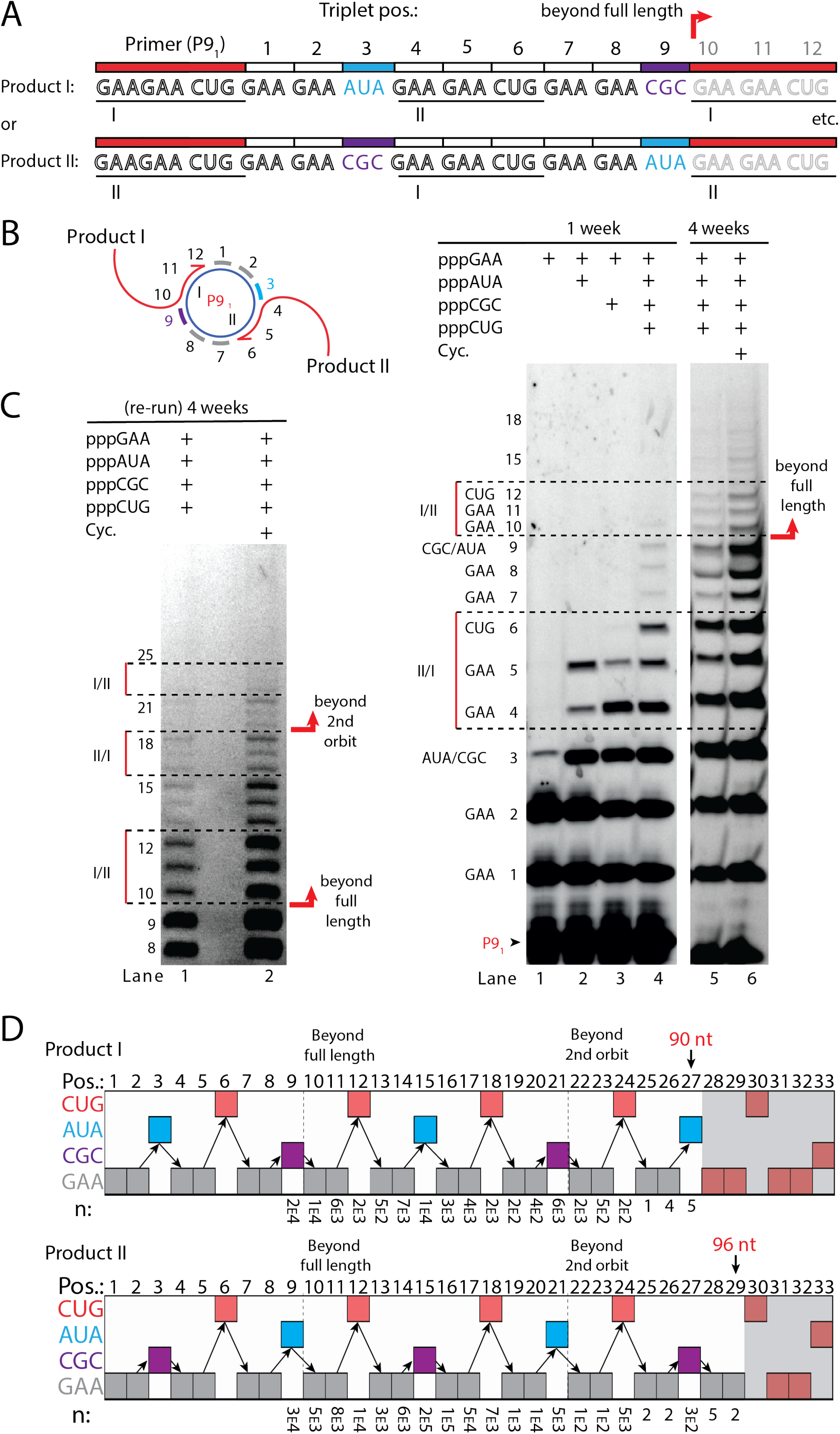
RNA-catalysed branched RCS. A) Product strand of primer extension experiments with scRNA template containing two priming sites (I and II) for primer P91. Depending on the priming site two different products will be made (I or II). B, C) Scheme and PAGE of primer extension experiments with only the noted triplets added with C) long electrophoretic separation to achieve optimal resolution of long products. Cycling (Cyc.) indicates that the samples had been exposed to four thermal and freeze-thaw cycles (80 °C 2 min, 17 °C 10 min, -70 °C 5-15 min, -7 °C 7 days) leading to increased efficiency. D) Sequencing of longer than full length branched RCS products on the double primer site scRNA (without cycling). Products I and II both reaching almost three full rounds of replication of the circular RNA template (up to 96 nt, 32 triplet incorporations). Original gel images and full analysis of the data in Figure 5D are supplied in Figure 5-source data 1.

Sequencing of the long, branched RCS RNA products (excised from band ≥15 triplets, Figure 5 - Figure supplement 1) identified a range of long reads (from both products I and II) including many reads of the product with 15 correct triplet incorporations (Figure 5E) representing ∼1.5 orbits (n: 7×10^3^ and 1×10^5^ reads of Product I and II, respectively). However, much longer sequences were present in decreasing numbers of reads, with the longest products comprising 29 correct triplet incorporations (96 nt) (n=2) representing RCS of more than 2.5 orbits and the longest reported product synthesised by the TPR. Thus, RNA-catalysed RCS has the potential to yield extended RNA concatemer products under isothermal conditions. Freeze-thaw (FT) cycles have been shown to enhance ribozyme activity by effecting RNA refolding (Mutschler et al., 2015) and indeed inclusion of 4 FT cycles lead to more efficient production of longer RCS RNA products (Figure 5B and C). In summary, isothermal conditions allow RCS of long concatemeric products containing multiple (>2.5) copies of the scRNA template with RCS efficiency further enhanced by FT cycling.

### Proto-viroid like self-circularizing ribozyme

A number of biological systems including viroids uses an RCS strategy for genome replication. However, RCS synthesis of RNA concatemers is only one part of the viroid replication cycle, which also involves resection (i.e. cleavage) of the concatemer into individual units and circularisation of unit length RNAs by ligation to recreate the original circular RNA genome. As both RNA cleavage and RNA ligation can be efficiently catalysed by RNA, we sought to investigate if a viroid-like replication cycle might be catalysed by RNA alone. To this end, we designed a proto-viroid RNA comprising a 39 nt scRNA template encoding a designed micro- hammerhead ribozyme (µHHz) as well as its substrate for cleavage (Figure 6A). The µHHz could be synthesized by the TPR (Figure 6B). Furthermore, the µHHz could catalyse both self-cleavage (forming a 2’,3’ cyclic phosphate (>p)) and re-ligation leading to circularization (under RCS reaction conditions at -7°C in eutectic ice) (Figure 6C, lane 2). A similar equilibrium between cleavage and ligation in eutectic ice had previously been observed for the unrelated hairpin ribozyme (Mutschler et al., 2015).

**Figure 6.**
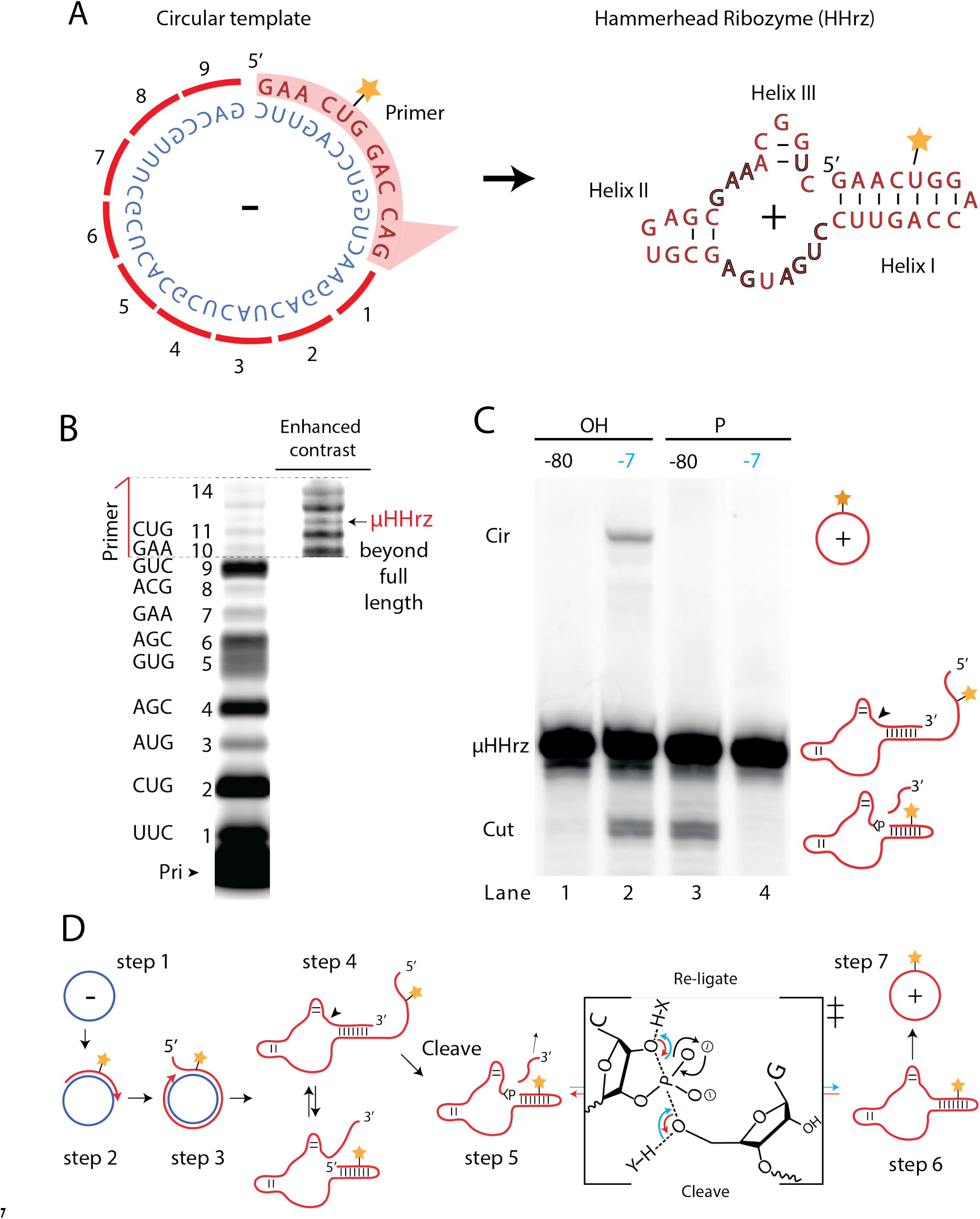
Steps of a viroid-like replication cycle catalysed by RNA alone. A) Illustration of the µHHrz (-) and (+) strand. B) PAGE gel showing primer extension of RCS synthesis of the µHHrz with substrate overhang to allow self-cleavage. C) PAGE gel showing cleavage and circularization of the µHHrz, but only when incubated at -7 °C, allowing eutectic phase to form, and with a free 5’-OH, needed for circularization, but not for cleavage. D) Schematic illustration of the RNA-catalysed viroid-like replication with steps comprising RNA-catalysed combined RCS (1-3), resection (4, 5) and self-circularization. (6, 7). Original gel images are supplied in Figure 6-source data 1.

When the µHHz had been 5’-phosphorylated (Figure 6C, lane 3) only cleavage but no circularization was seen, as phosphorylation blocks the 5’-hydroxyl nucleophile for re-ligation (see steps 5 and 6, Figure 6D). To the best of our knowledge, the µHHrz is the smallest (39 nt) self-cleaving and -circularizing RNA system reported to date and the first time self-circularization has been shown in a Hammerhead ribozyme. Kinetic analysis of the cleavage and circularization reaction show a slow but accumulating amount of cleavage product as a function of time (black points in Figure 6 - Figure supplement 1A). Analysing the ratio between linear (cleaved) and circular (ligated) products (Figure 6 - Figure supplement 1) showed that the proportion of circle was initially very high (approx. 40% after 0.5 day). Based on this, it is likely that all or most µHHz molecules are transiently circular at some point immediately after cleavage, but become progressively trapped in a state unable to re-ligate, most likely due to hydrolysis of the >p or misfolding. While only completing half of a full replication cycle (formation of a (+)-strand scRNA from a (-)-strand scRNA template, these results outline the potential for a full viroid-like rolling circle RNA replication cycle based on RNA-catalysis alone.

## Discussion

Viroids are transcriptional-parasites composed entirely of a circular RNA genome and are considered the simplest infectious pathogens known in nature. They lack protein coding regions in their genome, and can be completely replicated in ribosome-free conditions (Daròs et al., 1994; Diener, 2003; Fadda et al., 2003; Flores et al., 2009). They can comprise a circular RNA genome of as little as ∼300 nt in e.g. *Avsunviroidae* encoding a Hammerhead ribozyme responsible for maturation by resecting the RNA genome (Flores et al., 2000). The resected viroid genome is then ligated (circularized) by a host protein (e.g. tRNA ligase) (Nohales et al., 2012). Due to the simplicity of this replication strategy, viroids have been suggested to represent possible “relics” from a primordial RNA-based biology (Diener, 1989; Flores et al., 2014). Indeed circular RNA genomes would present a number of potential advantages for prebiotic RNA replication, including increased stability by end protection (Litke and Jaffrey, 2019), a reduced requirement for specific primer oligonucleotides to sustain replication (Attwater et al., 2018; Szostak, 2012) or resolve the end replication problem, i.e. the loss of genetic information from incomplete replication in linear genomes. Circular RNA structures self-assemble from RNA mononucleotides through wet-dry cycling (Hassenkam et al., 2020) and a virtual circular genome has been suggested as a model for primordial RNA replication (Zhou et al., 2021). Thus, Viroid-like systems are likely candidates to have emerged as the simplest Darwinian systems even before self-replication.

Here we have explored to what extent such a potentially prebiotic replication strategy can be carried out with RNA alone. Our data shows that RNA can indeed facilitate RCS on scRNA templates yielding concatemeric RNA products, which can be processed (i.e. resected) and recircularized by an encoded ribozyme in a scheme reminiscent of viroid replication. Thus, one half of full a viroid replication cycle ((-)-strand replication leading to a self-circularizing (+)- strand) can be carried out by just two ribozymes.

Completing the viroid replication cycle would require the reverse (+)-strand replication leading to a self-circularizing (-)-strand product and may require a second ribozyme (e.g. a second µHHrz) encoded in the (-)-strand akin to the mechanism used by natural viroids (Flores et al., 2000).

MD simulations indicate that the RCS process is aided by accumulating strain in the nascent dsRNA segment leading to increased peeling off of dsRNA 5’ and 3’ ends (i.e. strand displacement). In turn, this peeling off creates a more dynamic environment potentially aiding 5’ end invasion by extending the 3’ end. This topological strain induced strand displacement may be general and independent of the precise RCS mechanism on scRNA templates and thus should also apply to non-enzymatic polymerization of RNA. Our observation that RCS can be enhanced by the use of branched extension, freeze-thaw thermocycling and pre-freezing dilution may also relate to this. While the precise mechanistic basis for these enhancements is currently unknown, it seems plausible that all of these enhance 5’ product strand end displacement by accelerating conformational equilibration through RNA un- and refolding as observed previously (Mutschler et al., 2015).

In biology, both viroids and Hepatitis D virus (HDV) replication proceeds through RCS on circular RNA genomes mediated by proteinaceous RNA polymerases but RCS has also been reported for circular DNA templates and proteinaceous DNA polymerases in nature (Wawrzyniak et al., 2017) and in biotechnology (Daubendiek et al., 1995; Givskov et al., 2016; Kristoffersen et al., 2017; Mohsen and Kool, 2016). dsDNA persistence length is somewhat shorter than dsRNA (dsDNA: 45-50 nm (140-50 nt) vs. dsRNA 60 nm (200 nt) and stacking interactions weaker than in dsRNA (Kebbekus et al., 1995; Svozil et al., 2010), therefore dsDNA may more readily adopt a circular shape or kinks to alleviate build-up of strain or to adopt strong bends (Wolters and Wittig, 1989), we would nevertheless expect the a similar strand displacement effect would play part. Indeed, in both cases RCS proceeds efficiently for circular genomes ranging from a few hundred nt to over 1.5kb (Mohsen and Kool, 2016). In contrast, RNA-catalysed polymerization (record of producing approx. 200 nt products (Attwater et al., 2013)) is currently limited to RCS on small RNA circles. A more efficient RNA-catalysed RCS-based replication strategy will likely require improvements to the ribozyme polymerase catalytic activity, speed and processivity as well as the design of the template. Improvements to ribozyme polymerase processivity, which is known to be poor (Johnston et al., 2001; Lawrence and Bartel, 2003), might have the greatest impact and might be realized either through e.g. tethering or other topological linkages to the circular template (Cojocaru and Unrau, 2021). A more processive polymerase ribozyme should also result in less non-templated triplet TT activity, which appears to be a consequence of slow RCS extension and is likely aggravated by peeling off of the 3’ end. Thus more efficient RCS may also require the stabilization of the 3’ end triplet junction in the ribozyme active site in the same way as primer / nascent strand termini are stabilized within the active sites of proteinaceous RNA- and DNA polymerase (Chim et al., 2018; Houlihan et al., 2020). Finally, introduction of secondary structure motifs in the RNA nascent strand might drive increased 5’ dissociation (e.g. through formation of stable hairpin structures) relieving strain at the 3’-end.

Larger circular RNA templates might provide advantages for the RCS as they are less strained and provide increased access to the internal face of the circle and might also be able to encode the whole ribozyme itself. On the other hand, reduced torsional strain on the dsRNA would be expected to reduce strand invasion and “peeling off” of the product strand . All of these factors merit detailed investigation.

In conclusion, our motivation for investigating RNA-catalysed RCS was as a potential solution towards the so-called “strand inhibition problem”, the inhibition of RNA replication by exceedingly stable RNA duplex products. This inhibition has not just a thermodynamic component, i.e. the significant amount of energy required to melt such duplexes, but a kinetic component, because even if dissociation of RNA duplexes can be achieved, RNA replication must outpace duplex reannealing, which is rapid unless duplex concentrations are low or reactions take place in a highly viscuous medium (He et al., 2017; Tupper and Higgs, 2021). In this context, we reasoned that RCS might provide favourable properties: synthesis and strand displacement on a circular template can proceed essentially iso-energetically as base-pairing (H-bonding / stacking) interactions broken in the product strand during strand displacement are continuously compensated for by new base-pairing interactions formed in the nascent strand. In turn, this could lead to an open-end formation of template coupled stochiometric excess of single stranded RNA product strand to encode functions to further aid replication as we show here with resection and recircularisation by an encoded ribozyme.

In the course of this work, we discovered another mechanism that might contribute to overcoming the strand inhibition problem. MD simulations indicate that - at least - on small RNA circles – the build-up of strain in the nascent dsRNA could aid strand displacement (Figure 3). However, the MD simulations also suggest that strain is non-directional destabilizes both nascent strand 5’- as well as 3’-end likely inhibiting extension and promoting non-templated triplet addition. Thus, the potential advantages of scRNA RCS seems to be tempered by opposing effects such as strain as well as reduced template accessibility due to circular RNA ring geometry (Figure 1). Nevertheless, we find that a viroid-like replication strategy can be accomplished by RNA catalysis alone, with one ribozyme performing RCS on circular RNA templates yielding concatemeric RNA products, which can be processed (i.e. resected) and recircularized by a second ribozyme. Future improvements in polymerase ribozyme activity and processivity may allow all necessary components of such a replication cycle to be encoded on a circular RNA “genome” and propagated by self-replication and - processing reactions.

## Materials and methods

### Oligonucleotides

Base sequences of all oligonucleotides used throughout this work can be found in Supplementary file 1.

### In vitro transcription

dsDNA templates (containing T7 promotor sequence at the 5’ end upstream of the region to transcribe) for *in vitro* transcription was generated by “fill-in” using three cycles of mutual extension using GoTaq HotStart, (Promega, Madison, Wisconsin) between the relevant oligonucleotide and primers: 5T7 or HDVrt (the latter for defined 3’ terminus formation (Schürer et al., 2002))

The T7 transcription protocol used is based on Megascript. Briefly explained, transcriptions of RNA requiring a triphosphate at the 5’-end (termed GTP Transcription) reaction were carried out under the following conditions: 40 mM Tris-Cl pH 8, 10 mM DTT, 2 mM spermidine, 20 mM MgCl2, 7.5 mM each NTP (Thermo Fisher Scientific), dsDNA templates (varying amount, preferably >5 μM), 0.01 units/μL of inorganic pyrophosphatase (Thermo Fisher Scientific, Waltham, Massachusetts), ∼50 μg/μL of T7 RNA polymerase (home made by Isaac Gallego). Reactions were left overnight (∼16 hours) at 37°C. Then 0.5x volume EDTA (0.5 M) was added together with (at least) 2.5x volume of loading buffer (final conditions >50% formamide or >8M Urea and 5 mM EDTA). For transcription of RNA with a monophosphate 5’-end (termed GMP Transcription) the same procedure is followed as for NTP Transcription, however 10 mM GMP and 2.5 mM of each NTP instead of the higher amount of NTP used for GTP transcription.

### Gel electroporation for analysis or purification

The sample in appropriate loading buffer were separated on 10-20% 8 M Urea denaturing PAGE gel using an EV400 DNA Sequencing Unit (Cambridge Electrophoresis). The product band was visualised by UV shadowing (for non-labelled RNA) or florescence scanning (Typhoon scanner, Amersham Typhoon) (for labelled RNA). When needed the identified product based on relative migration was excised. The excised gel fragment was then thoroughly crushed using a pipette tip and suspended in 10 mM Tris-Cl pH 7.4 to form a slurry. For freeze and squeeze extraction, the slurry was frozen in dry ice, then heated to 50°C (∼5 min) and finally left rotating at room temperature (from 2 hours to overnight) to elute the product from the gel material. The eluate was then filtered using a Spin-X column (0.22 μm pore size, Costar), ethanol precipitated, (100 mM Acidic acid and 80% ethanol (10 ug glycogen carrier was present when noted)). UV absorbance was measured with a Nanodrop ND-1000 spectrophotometer (Thermo Fisher Scientific) to determine yield of redissolved purified RNA.

### Calculation of extension efficiency

Gel Images from the Typhoon scanner where analysed in ImageQuant software (Cytiva life science) for quantifying band-intensity. Quantified band intensities were exported to Excel (Microsoft, Redmond, Washington) for further analysis. Extension efficiency (*E*) for a given band (*b*) was calculated as the sum of the intensities (*I*) of all the bands from *b* to *n*, (*n* being the highest detectable band), divided by the sum of *I* of all bands from *b-*1 to *n*:

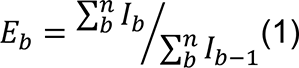

Thus, *E* represents the efficiency of the given ligation junction (Lb) to allow production of the extension product in band *b, i.e.* the extension efficiency.

### Triplet transcription

Triplets were prepared via run-off in vitro transcription with T7 RNA polymerase. More details on the method can be found in (Attwater et al., 2018). Reaction conditions were as follows: 100 pmols of DNA template for each triplet was mixed with equimolar DNA oligo 5T7 to form the template for transcription. For triplets starting with purines, the NTP transcription protocol was used as described above with a total NTP concentration of 30 mM but only adding the nucleotides necessary for the triplet (e.g. AUA was transcribed with only ATP and UTP). For triplets starting in pyrimidines a lower total NTP concentration was used (4.32 mM) as this yielded better defined bands for purification. 50 μL transcription reactions were stopped with 2 μL EDTA (0.5M) and 5 μL of 100% glycerol was added to facilitate gel loading. The samples were separated by 30% 3 M Urea denaturing PAGE as described above. Correct sequence composition was confirmed by A260/280 absorbance ratio, measured with the Nanodrop.

### Circularization of RNA

Linear 5’-end monophosphate labelled RNA to be used for circularization was either prepared by *in vitro* transcription (300 μL reaction volume) or ordered directly as chemically synthesized RNA (Integrated DNA Technologies (IDT), Iowa, United Stated). Linear RNA was gel purified as described above. When needed purified RNA was treated with T4 polynucleotide kinase (PNK) (New England Biolabs (NEB), Ipswich, Massachusetts) to remove 3’-end cyclic phosphate then RNA was phenol/chloroform extracted, ethanol precipitated and redissolved in ddH2O. For splinted ligation, 3 pmol purified RNA was mixed with equimolar splint RNA in 262.5 μL ddH2O. The sample was heated to 80°C (2 min.) followed by cooling to 17°C (10 min.) and finally incubated on ice (5-30 min.). Then reaction conditions were adjusted to 50 mM Tris-HCl pH 7.5, 2 mM MgCl2, 1 mM DTT, 400 μM ATP (1x T4 RNA ligase 2 reaction buffer (NEB)) including 0.25 units/μL T4 RNA ligase 2 (Neb) (final volume 300 μL) and samples left over night (∼16 hours) at 4°C. For non-splinted ligation, 10 pmol gel purified RNA was mixed in 237 μL ddH2O followed by heating to 95°C and then quickly moved to ice. Then reaction conditions were adjusted to 50 mM Tris-HCl, pH 7.5, 10 mM MgCl2, 1 mM DTT (1x T4 RNA ligase reaction buffer (Neb)), 100 μM ATP including 1 unit/μL T4 RNA ligase 1 (NEB) (final volume 300 μL) and samples left over night (∼16 hours) at 16°C. Circularized RNA was electrophorated by 10% 8M Urea denaturing PAGE for analysis and purification as described above.

### Templated RNA-catalysed RNA synthesis (the primer extension assay)

Ribozyme activity assay was performed essentially as described in (Attwater et al., 2018). In a standard reaction (modified where specified), ribozyme heterodimer (5TU/t1), template, primer (5 pmol of each) and triplets (50 pmol of each) were annealed in 7.5 μL water (80 °C 2 min, 17 °C 10 min). Then 2.5 μL 4x reaction buffer was added (final volume 10 μL) and samples were left on ice for ∼5 min to ensure folding. Final pre-frozen conditions were (unless otherwise noted) either (Tris buffer system) 50 mM Tris (pH 8.3 at 25 °C), 100 mM MgCl2 and 0.01% Tween20, or (CHES buffer system) 50 mM CHES (pH 9 at 25 °C), 150 mM KCl, 10 mM MgCl2 and 0.01% Tween20. At this point some samples (noted in the text) were diluted by adding ddH2O (e.g. 50x dilution corresponds to adding 490 μL ddH2O to the 10 μL samples). Finally, samples were frozen on dry ice and incubated at -7 °C in a R4 series TC120 refrigerated cooling bath (Grant,Shepreth, UK) to allow eutectic phase formation and reaction, respectively.

To end the incubations, samples that had been diluted were thawed, moved to 2mL tubes, ethanol precipitated (with glycogen carrier) and redissovled in 10 μL ddH2O. This step was avoided for undiluted samples that were already 10 μL. Finally, 0.5 μL EDTA (0.5M) was added to all samples to a final volume of 15 μL. (In experiments where the effect of dilution was investigated, e.g. as experiment presented in Supporting Figure 5, ddH2O was added to all the thawed samples to reach the same volume before precipitation).

To prepare for separation of extension products, 3 μL of the reacted samples after addition of EDTA (corresponding to 1 pmol template RNA) was diluted to reach the final loading conditions: 166 mM EDTA, 6M Urea (+ Bromophenol blue) and 10-20 pmol competing RNA (to prevent long product/template reannealing) (final volume 10 μL). Finally, samples were denatured (95°C for 5 min) and RNA separated by 8M Urea denaturing PAGE.

### Sequencing of extension products

In the primer extension reactions used for sequencing, the primer extension was performed as described above except for the following changes: 5 pmol ribozyme heterodimer/template, 20 pmol primer (with a 5’adapter sequence) and 100 pmol of each triplet was used. In the cases where multiple templates were mixed in the same reaction (one-pot), the final template concentration remained 5 pmol in total. All reactions were done in the CHES buffer and were diluted 50x as standard.

#### Adapter ligation and RT-PCR

After Urea PAGE separation of the extension products, the noted region of the gel was dissected out, and carefully recovered as described above. The RNA was ethanol precipitated (80% ethanol with 10 ug glycogen carrier) resulting in a dry RNA pellet. To append an adaptor sequence to the 3’-end of the purified RNA products the dry RNA was redissolved in conditions allowing adenylated adapter ligation by T4 RNA Ligase 2 truncated K227Q (Neb) following manufacturers descriptions. Final adapter ligation conditions were: 50 mM Tris-HCl, pH 7.5, 10 mM MgCl2, 1 mM DTT (1x T4 RNA ligase reaction buffer (NEB)), 15 % PEG8000, 0.04% Tween 20, 5 pmol adenylated DNA primer (Adap1, for base sequences see Supplementary file 1) and 20 U/µL T4 RNA Ligase 2 truncated K227Q (Neb) (final volume 10 µL). The samples were then ligated at 16°C for two hours. Pre-adenylation of Adap1 using Mth RNA Ligase (Neb) was performed following manufacturers descriptions. After adapter ligation, samples were diluted 10-fold to achieve conditions for performing RT-PCR (25 cycles) using 0.5 µM forward (PCRp3) and reverse primer (RTp1) and the SuperScript III One-Step RT-PCR system with Platinum Taq DNA polymerase (Thermo Fisher Scientific). Finally, RT-PCR products were gel purified in 3% agarose gel and cleaned up using QIAGen gel extraction kit (QiaGen, Hilden, Germany).

#### Sanger sequencing

Purified RT-PCR products were cloned in to pGEM vector using pGEM-T easy vector Systems (Promega) as described by manufacturer and transformed into heat- competent 10-Beta cells (NEB). Inserts from single colonies were PCR amplified (using primers pGEM_T7_Fo, pGEM_SP6_Ba) and send in for Sanger sequencing (Source bioscience) (using pGEM_T7_Fo as sequencing primer). Illumina sequencing: Illumina adaptors were added to purified RT-PCR products by PCR (15 cycles) using 0.5 µM forward (Ill*x*_Fo, *x* denotes different barcodes 1-15, see oligo sequences in supplementary material) and reverse primers (Ill_Ba) and Q5 Hot-Start High-Fidelity 2X Master Mix (Neb). PCR products were gel purified in 3% agarose gel and qPCRed (using NEBNext Library Quant Kit for Illumina) to quantify concentration. Finally, the DNA (consisting of Illumina adapters, barcodes and RT-PCRed sequence from the RNA extension) were prepared following manufactures protocol for MiSeq Illumina sequencing (Illumina, San Diego, California) (see e.g. MiSeq System Guide).

### Sequencing data analysis

Illumina Sequencing data were acquired and processed as FASTQ files using Terminal (and available software packages such as FASTX-toolkit). Prior to analysis the whole output file form illumina sequencing runs (containing also unrelated sequences) was split based on barcodes identifying the individual samples and trimmed starting with the original (P9_1_) primer sequence (GAAGAACTG). After the P9_1_ sequence, the triplets at positions 1, 2 ,3 etc. would be identifiable representing extension products made by the ribozyme. The presence for the 3’ adapter sequence (GTCGAATAT…) in the aligned sequences marked the end of the original RNA extension product. Sequencing data can be found as described belos under section data availability: File 1 include sequence data for circular and linear one-pot analysis (C1 and L1, respectively), File 2 include sequence data for branched RCS analysis (B3)). *Analysis of the one-pot experiments:* By counting the number of times a given triplet was present at a given position, we were able to calculate the fidelity for each triplet at this position. Identifying and counting the sequencing reads (n) for each position was done using *grep* (in Terminal) with a list of all relevant sequences (positions 3 to 18) and the sequencing files. The triplet at position 3, the first barcode position, was used to classify the sequences into coming from template A to D and thus has 100% fidelity for the correct triplet (Figure 4B).

In example, for analysing the fidelity (*F*) of position 4, the following list was used: GAAGAACTG_(primer)_GAA_(pos1)_GAA_(pos2)_YYY_(pos3)_XXX_(pos4)_. Here YYY was either of the first barcode triplets for templates A-D, (ATA_(template A)_, AAA_(template B)_, TTA_(template C)_ or ATC_(template D)_) and XXX was either of the 14 possible triplets (CTG, ATA, CCA, CCC, AAA, CAC, GGG, TTA, TCC, GGC, ATC, GAT, CGC, GAA). *F* at position 4 was then calculated for template A-D as the number of occurrences of a triplet in positon 4 (e.g. CCA) divided by the sum of occurrences of all the triplets multiplied by 100%. A generalized term for calculating the *F* at all positions (3-18) and for all templates (A-D) is:

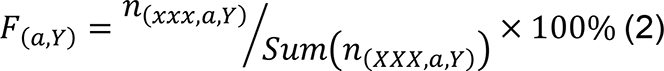

Here *F* is the fidelity, *a* is the position of the triplet, *Y* is the template A-D. *n* is the number of sequencing reads for a given triplet (xxx) for position *a* on template *Y* or for all the 14 triplets (XXX), for *a* on *Y.* Eventually, the fidelity for positions 3-15 in the context of template A-D for all triplets was plotted in Figure 4B. Accumulated chance for a product of reaching positions X (shown in plot in Figure 4C) was calculated by multiplying all fidelities for moving from position 3 to position X with correct triplets (fidelities found in Figure 4C). Data for this analysis can be found as described in data availability section below (File 1). Numerical data and calculation is supplied in Figure 4-source data 1.

*Analysis of the branched RCS:* By counting the number (n) of correct sequences with a specific length ending in the 3’adapter sequence, we identified long RCS products (Figure 5D). This was done using *grep* (in Terminal) with a list of all relevant sequences (positions 9 to 30, both product I and II), and the sequencing file. Data for this analysis can be found as described in data availability section below (File 2) Numerical data and calculation is supplied in Figure 5-source data 1.

### Self-circularizing Micro Hammerhead ribozyme assay

RNA catalysed synthesis of fluorophore labelled self-circularizing micro Hammerhead ribozyme was prepared in 2x large (500 pmol) reactions set up and incubated as described above. Specifically, 500pmol ribozyme heterodimer (5TU/t1) and circular template (HHrzCtemp alt7), 2000 pmol primer (HHrzP12) and 50 µmol of each of the triplets were annealed followed by adding buffer to 50 mM CHES, pH 9; 150 mM KCl; 10 mM MgCl2, 0.05% Tween 20 (1mL). Then the sample was diluted 50 times to a final volume of 50 mL. After 4 weeks incubation at -7 °C, EDTA was added (5mM final concentration), reactions were thawed and concentrated to a final volume of ∼300 µL using a centrifugation filter (Amicon Ultra, 3 kDa cut off) retaining long RNA products. Micro Hammerhead ribozyme RNA (marked in Figure 6C) was purified by gel electrophoresis and excised product was dissolved to 10 µM in H2O with 0.5 mM EDTA.

Chemically synthesized fluorophore labelled self-circularizing micro Hammerhead ribozyme RNA (IDT) was gel purified as described above and excised product was dissolved to 10 µM in H2O with 0.5mM EDTA.

### micro Hammerhead ribozyme cleavage/circularization assay

Micro hammerhead self-circularization assays comprise 10 pmol micro HHrz annealed (80 °C 2 min, 17 °C 10 min) in 4 µL water with 1 µL 5x reaction buffer, final reaction conditions: 50 mM CHES, pH 9; 150 mM KCl; 10 mM MgCl2 (same as for the Templated RNA-catalysed RNA synthesis). Then incubated in ice for 5 min to ensure folding. This was then frozen on dry ice and either moved to -7 °C for eutectic phase formation (reaction) or -80 °C (control). After incubation, 10 µL loading buffer (95% Formamide, 25 mM EDTA, Bromophenol blue) was added directly to the cold samples to stop the reaction and mixed while thawing. Finally, reactions were analysed by 20% denaturing PAGE like described above.

5’ phosphorylation of micro HHrz RNA with polynucleotide kinase (NEB) as done following manufacturer’s directions. RNA was then phenol/chloroform washed, precipitate and dissolved in ddH2O with 0.5mM EDTA to 10 µM (determined by Nanodrop).

### Molecular Dynamics simulations

All simulations were set up with the AMBER 18 suite of programs and performed using the CUDA implementation of AMBER’s pmemd program (Case, n.d.). A linear ssRNA of 36 nt with the sequence (UUC)_12_ was built using the NAB utility, which was then circularised using an in-house programme (Pyne et al., 2021). From there, the complementary strand containing GAA triplets was progressively grown representing the different stages of the rolling circle replication, containing 9, 18, 21, 24, 27 till 30 nt of dsRNA keeping the rest single-stranded. For each stage, a representative structure was used as a scaffold to grow the dsRNA part and thus build the structure to model next stage. A linear dsRNA fragment containing 4 GAA triplets with a nick between the first and second was run as a control. This molecule had a total length of 16 bp as it was capped by a CG dimer on each end.

The AMBER99 forcefield (Cheatham et al., 1999) with different corrections for backbone dihedral angles including the parmBSC0 for α and γ (Pérez et al., 2007) and the parmOL3 for χ (glycosidic bond) (Zgarbová et al., 2011) were used to describe the RNA. All initial structures were explicitly solvated using a truncated octahedral TIP3P box with a 14 Å buffer. They were neutralized by two different types of salt, KCl and MgCl_2_, described by the ‘scaled charged’ Empirical Continuum Correction (ECC) set of ion parameters (Duboué-Dijon et al., 2018), and with the necessary ion pairs (Machado and Pantano, 2020) for matching 0.2 M in the case of KCl, and 0.1 and 0.5 M in the case of MgCl_2_. Simulations were performed at constant T and P (300 K and 1 atm) following standard protocols (Noy and Golestanian, 2010) for 400 ns.

The last 100 ns sampled every 10 ps were used for the subsequent analysis. AMBER program CPPTRAJ (Roe and Cheatham, 2013) was used to determine base-pair step parameters, radial distribution functions of ions around RNA and distances between atoms, including groove width and hydrogen bonds. The latter were defined with a distance cutoff of 3.5 Å and an angle cutoff of 120°. Counterion-density maps were obtained using Canion (Lavery et al., 2014) and were subsequently visualized with Chimera (Pettersen et al., 2004). SerraNA software was used to calculate curvatures at different sub-fragment lengths (Velasco-Berrelleza et al., 2020).

## Acknowledgements

This work was supported by the Carlsberg Foundation (ELK), by the Medical Research Council (MRC) program grant program no. MC_U105178804 (PH), by the Engineering and Physical Sciences Research Council (EPSRC) grant EP/N027639/1 (AN) and by the EPSRC (EP/R513386/1) (MB). Simulations were performed on JADE (EP/T022205/1)

Thanks to the HecBiosim consortium (EP/R029407/1), Cambridge Tier-2 (EP/P020259/1) and the local York facilities.

## Author contributions

ELK and PH conceived and designed experiments. ELK performed all experiments except molecular dynamics simulation (AN). All authors analysed data, discussed results and co-wrote the manuscript.

## Additional files

Supplementary file 1. Oligonucleotide sequences.

Transparent reporting form.

## Data availability

Simulations are available at the University of York Data Repository (<u>10.15124/b92977bd-f016-4740-8b4a-f86c68d5eb2c</u>).

Sequencing data used for analysis presented in Figure 4 (File 1) and 5 (File 2) are available on Dryad (https://doi.org/10.5061/dryad.tht76hf10).

### Figure supplements and Movies

**Figure 1 - Figure supplement 1.**
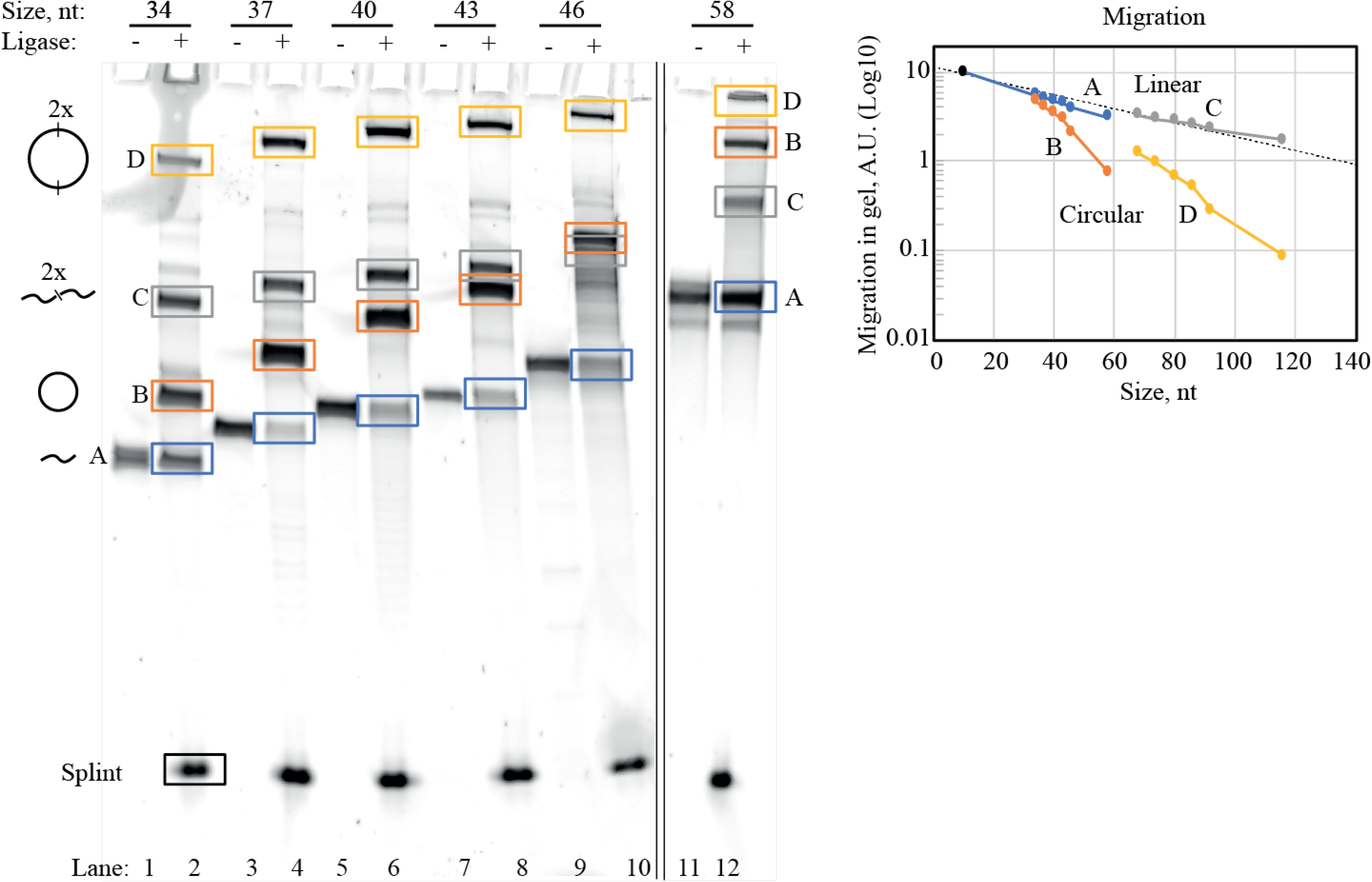
Purification of circularized RNA. Identification and dissection of circularized RNA were performed by denaturing PAGE. Representative SyBr Gold stained 10% Urea PAGE gel is shown here for illustrating the circularization process of circular RNA templates used in main text Figure 2B. In the gel, RNA before (odd lanes) and after (even lanes) ligation with T4RNA ligase 2 was analysed. A 10 nucleotide (nt) RNA splint (covering 5 nt of the 5’-end and 3’-end of the linear RNA strand) was used for circularization as required by T4 RNA ligase 2. In the ligated samples, multiple bands (A-D) appeared representing various combinations of ligated RNA strands. By migration analysis (right panel), we identified A and C as linear constructs and B and D as circularized constructs (illustrations of the identified structure of A-D can be seen to the right of the gel). Monomeric circularized RNA (corresponding to band B) was dissected out and used throughout this work. Bands A-D discussed here should not be confused with templates A-D used in main text figure 4. Original gel is supplied in Figure 1-Figure supplement 1-source data 1.

**Figure 2 - Figure supplement 1.**
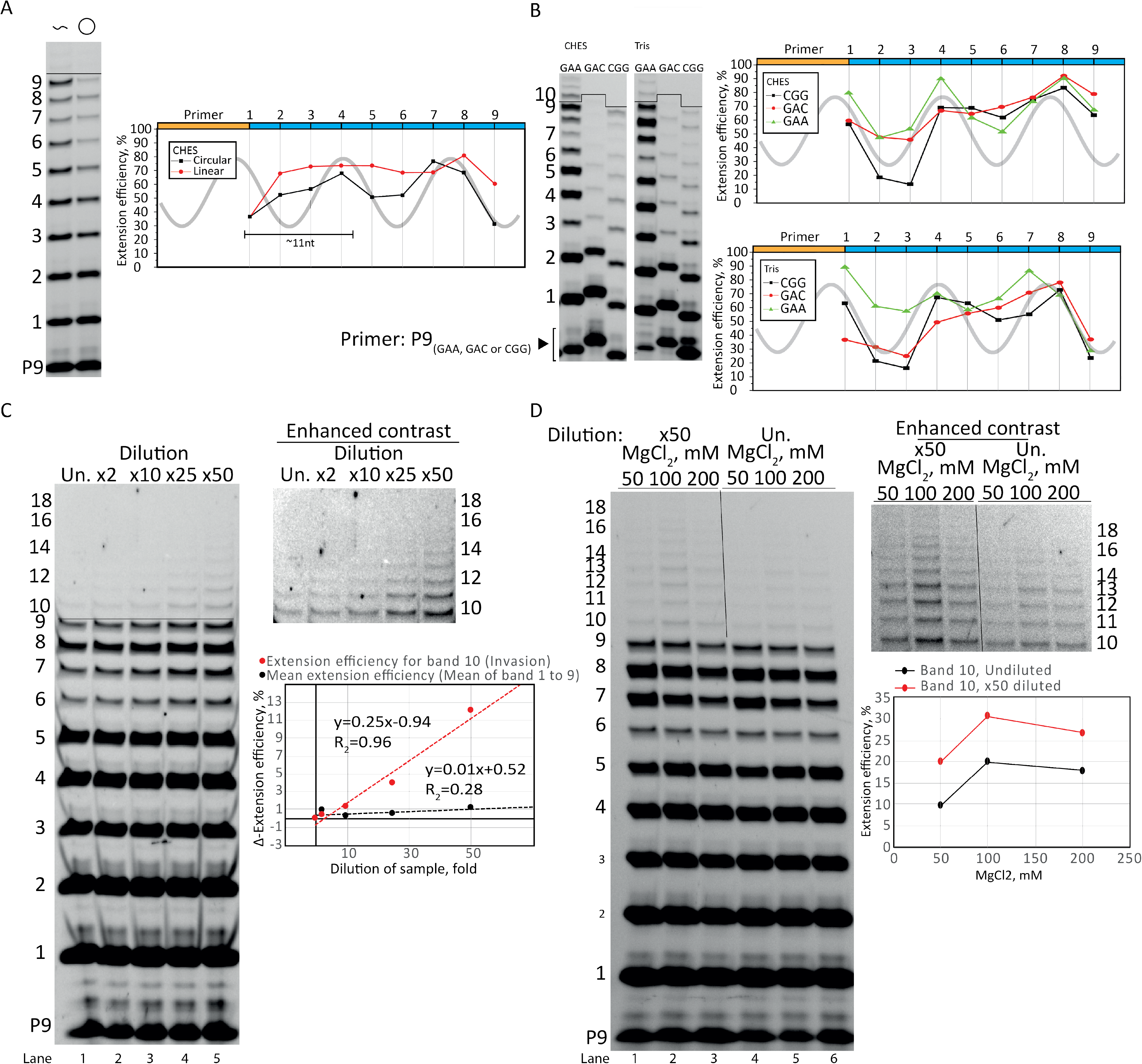
Optimization of Rolling Circle Synthesis. A) Comparison between linear and circular primer extension using the CHES reaction buffer system (similar to main text Figure 1D that is in the Tris buffer system). Extensions performed at -7 °C for 2 weeks. B) The periodic oscillations were observed with various repeat sequence templates (CGG, GAC and GAA) in both CHES and Tris buffer systems. Extensions performed at -7 °C for 4 weeks. C) Dilution of samples increased the efficiency. The plot in C) shows the difference in extension efficiency (Δ-extension efficiency) between the undiluted (Un.) and the 2-50 fold diluted (x2-x50) samples. The Δ-extension efficient of the ligations before invasion (mean of bands 1-9) were unaffected by the dilution (giving a Δ-extension efficient of ∼1). However, the Δ-extension efficiency of band 10 (full length +1 triplet, invasion) increased strongly with dilution. Extensions performed at -7 °C for 1 week. D) The same effect of dilution (improving invasion) was seen over a range of MgCl2 concentrations (50-200 mM). Extensions performed at -7 °C for 1 weeks. All extension reactions presented here were run at standard reaction conditions described in main text Materials and Methods except when specified otherwise for dilution, salt or buffer system. E) Image of the whole gel where parts are shown in main text figure 2D. Original gels and numeric values are supplied in Figure 2-Figure supplement 1-source data 1.

**Figure 3 - Figure supplement 1.**
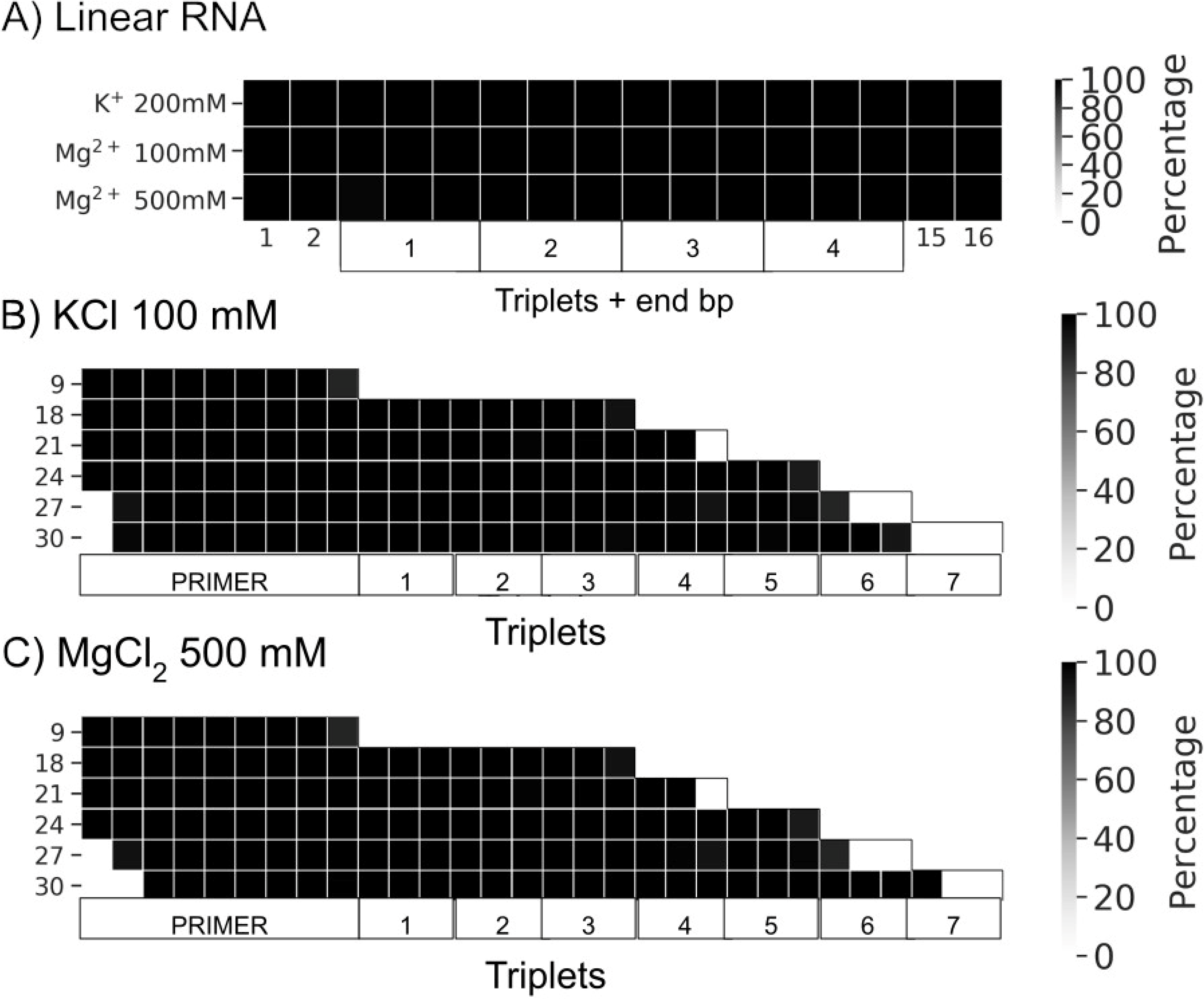
Percentage of frames from the last 100 ns of the simulations presenting canonical hydrogen bond pairing for each base pare (bp): A) Linear RNAs solvated with the three buffers (100 mM KCl, 200 mM MgCl_2_ and 500 mM MgCl_2_); B) Rolling circle RNA synthesis (RCS) simulations solvated with 100 mM KCl; C) RCS simulations solvated with 500 mM Mg Cl_2_.

**Figure 3 - Figure supplement 2.**
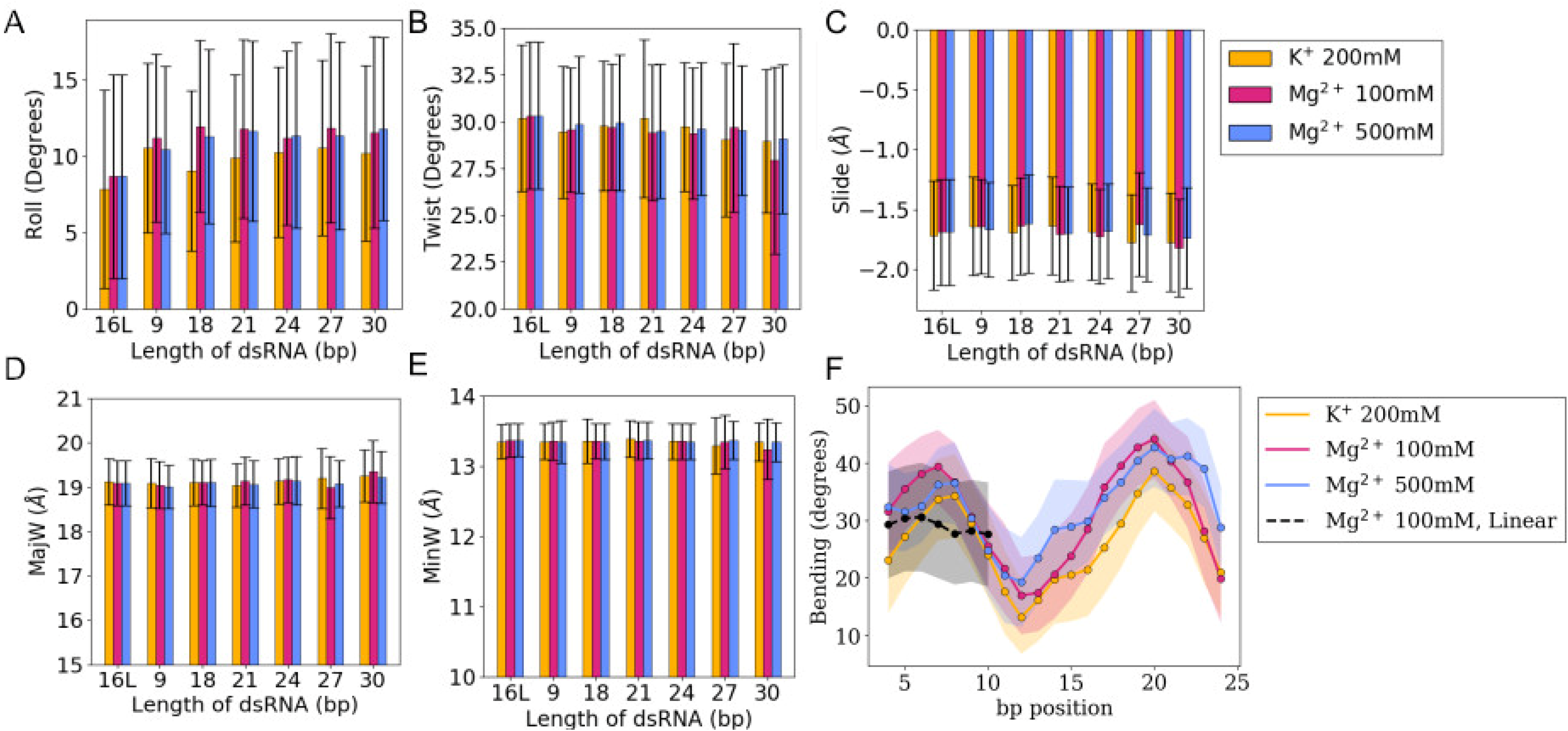
A-E) Averages and standard deviations (as error bars) of bp-step parameters (roll, slide and twist) together with major and minor groove widths (MajW and MinW, respectively) calculated over the last 100 ns of the simulations. The trajectory of the 16 bp linear RNA is labelled as 16L. F) Bending profile for all the sub-fragments 4 bp-long along 30 bp of dsRNA embedded in a 36-bp circular ssRNA.

**Figure 3 - Figure supplement 3.**
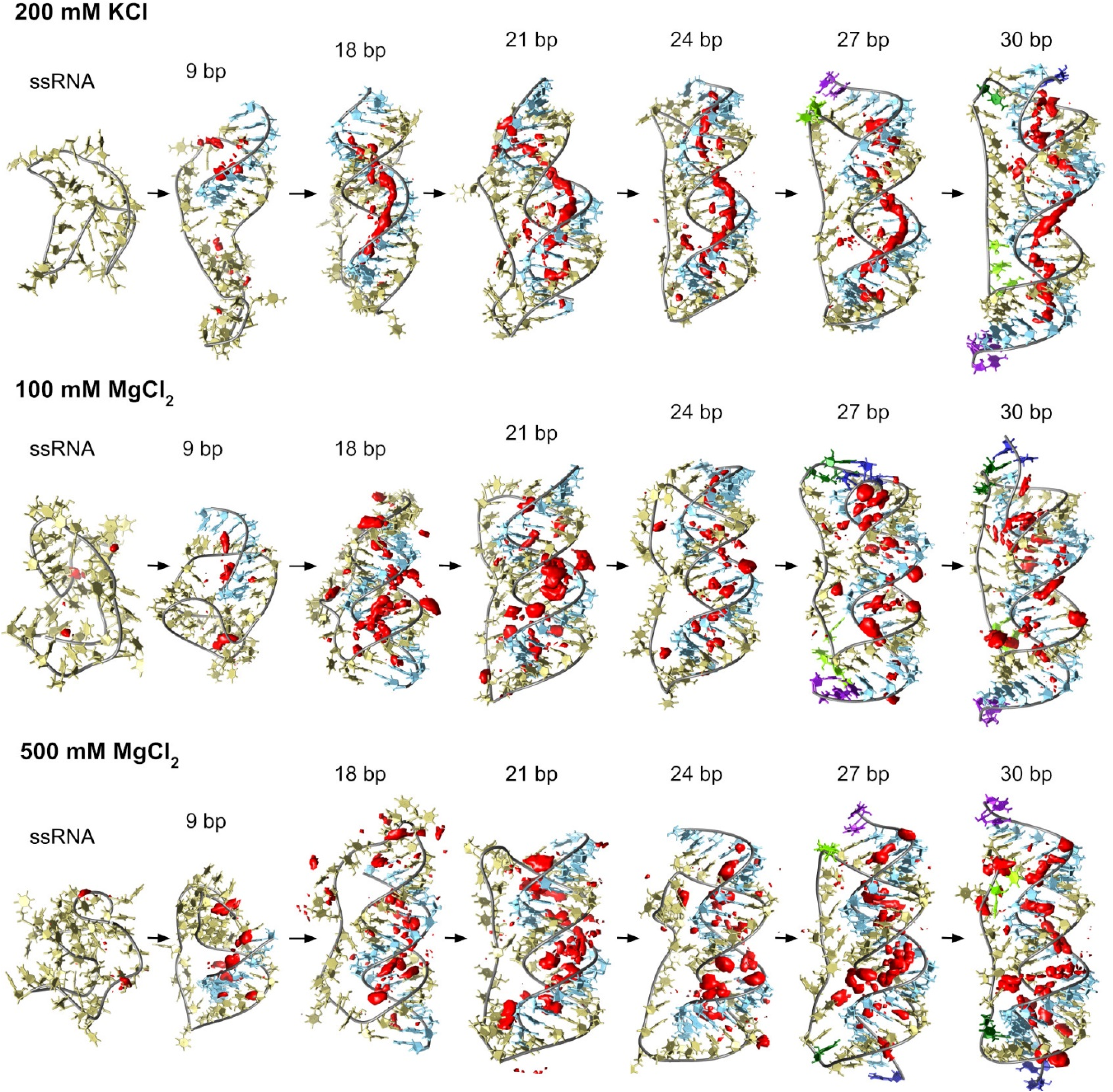
Counterion-density maps around RNA molecules that show an occupancy ∼10 times or greater the bulk concentration (in red) as seen in simulations. These areas are the molecular regions where cations localize preferentially. In the case of 200 mM KCl, these align along the grooves, whereas, in the case of MgCl_2_, they tend to be closer to the backbone and bridge distant backbone points, making the bases more exposed. Extremely high Mg^2+^ concentrations provide similar interacting profiles to moderate levels indicating saturation on the preferred binding sites in both cases.

**Figure 3 - Figure supplement 4.**
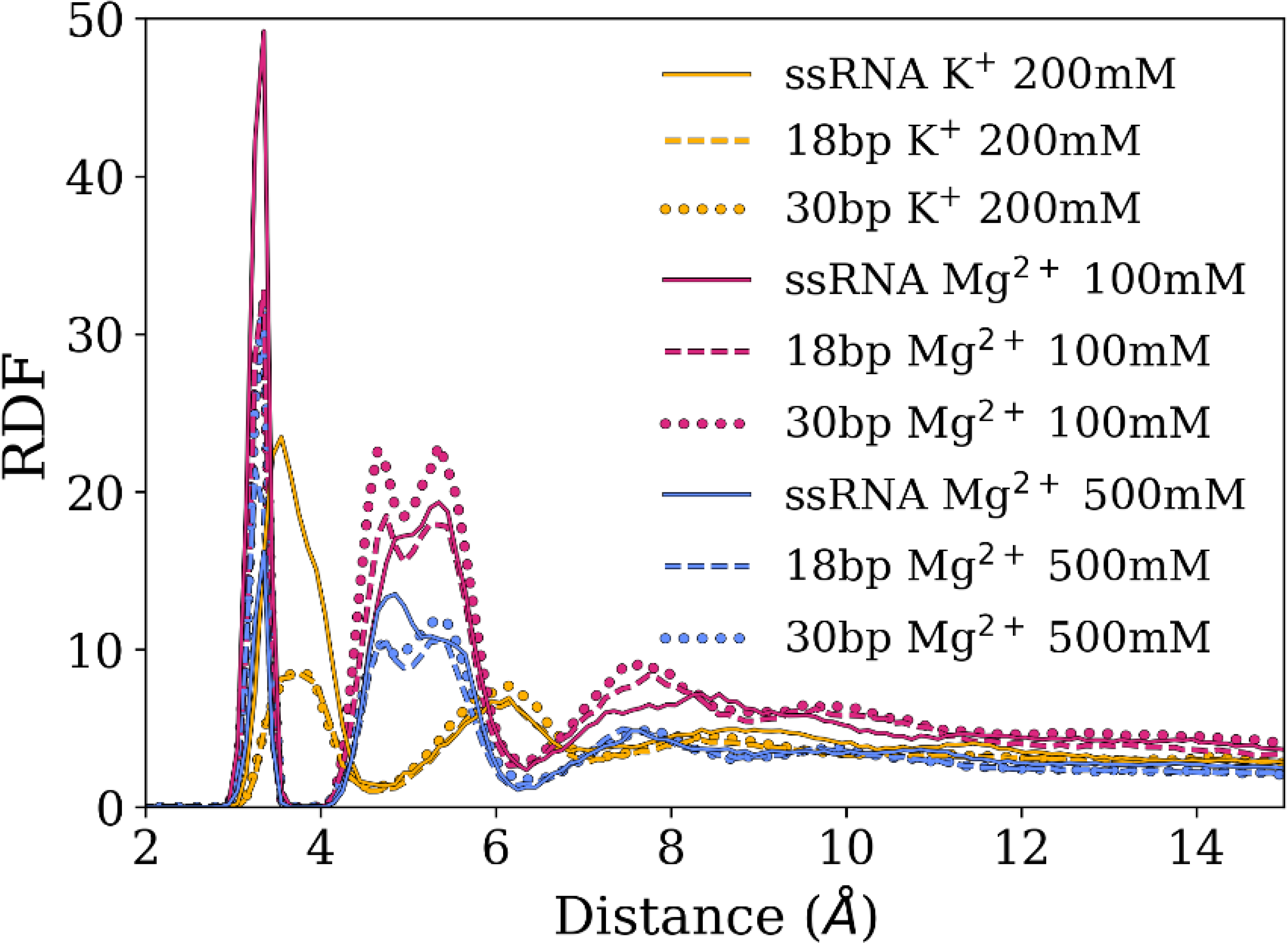
Averages and standard deviations (as error bars) of radial distribution functions (rdf) of cations around RNA backbone phosphates. The rdf indicate the probability of finding an ion within a certain distance of a particular RNA atom in relation to its bulk concentration (set at 1). Magnesium ions make more direct interactions with RNA backbone (first peak) and mediated by water molecules (subsequent peaks) than potassium. The smaller rdf peaks observed on 500 mM compared to 100 mM indicate a relatively lower ion condensation around RNA with respect to bulk concentration due to saturation.

**Figure 4 - Figure supplement 1.**
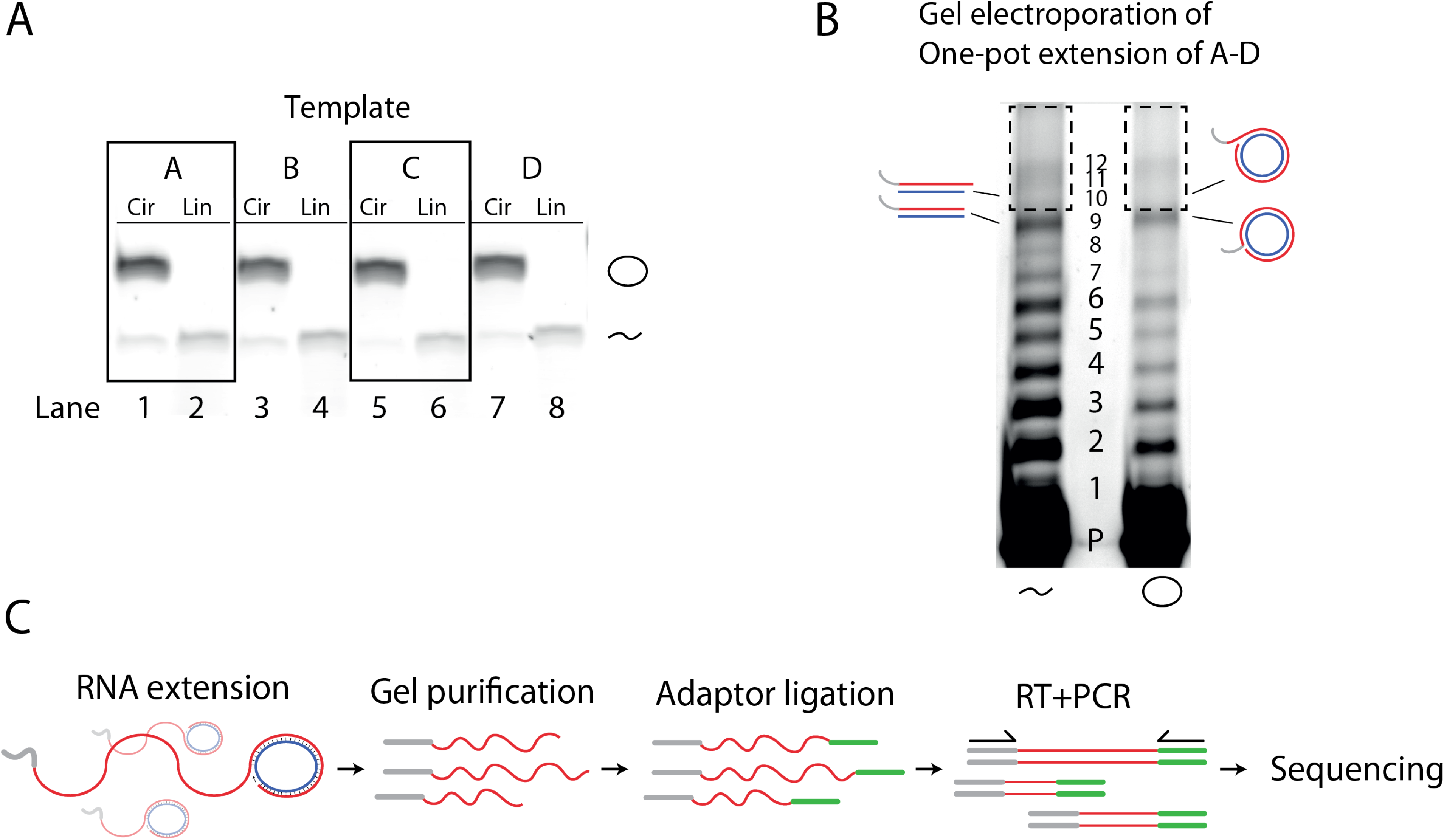
Deep sequencing of extension products. A) Representative SyBr Gold stained 10% Urea PAGE gel showing linear and circularized circular templates A-D. B) 10% Urea PAGE separation of one-pot extension reaction used for deep sequencing. Dashed box denotes excised region (above the full-length product (band 9)) used for RNA recovery and Deep-sequencing. C) Illustration of the protocol for sequencing of extension products. The initial extension products gets gel purified, then 3’-adaptor ligated with a 5’- adynalated DNA adapter strand, and finally RT-PCR amplified (adding additional adapter sequences) and submitted for sequencing. Original gels are supplied in Figure 4-Figure supplement 1-source data 1.

**Figure 4 - Figure supplement 2.**
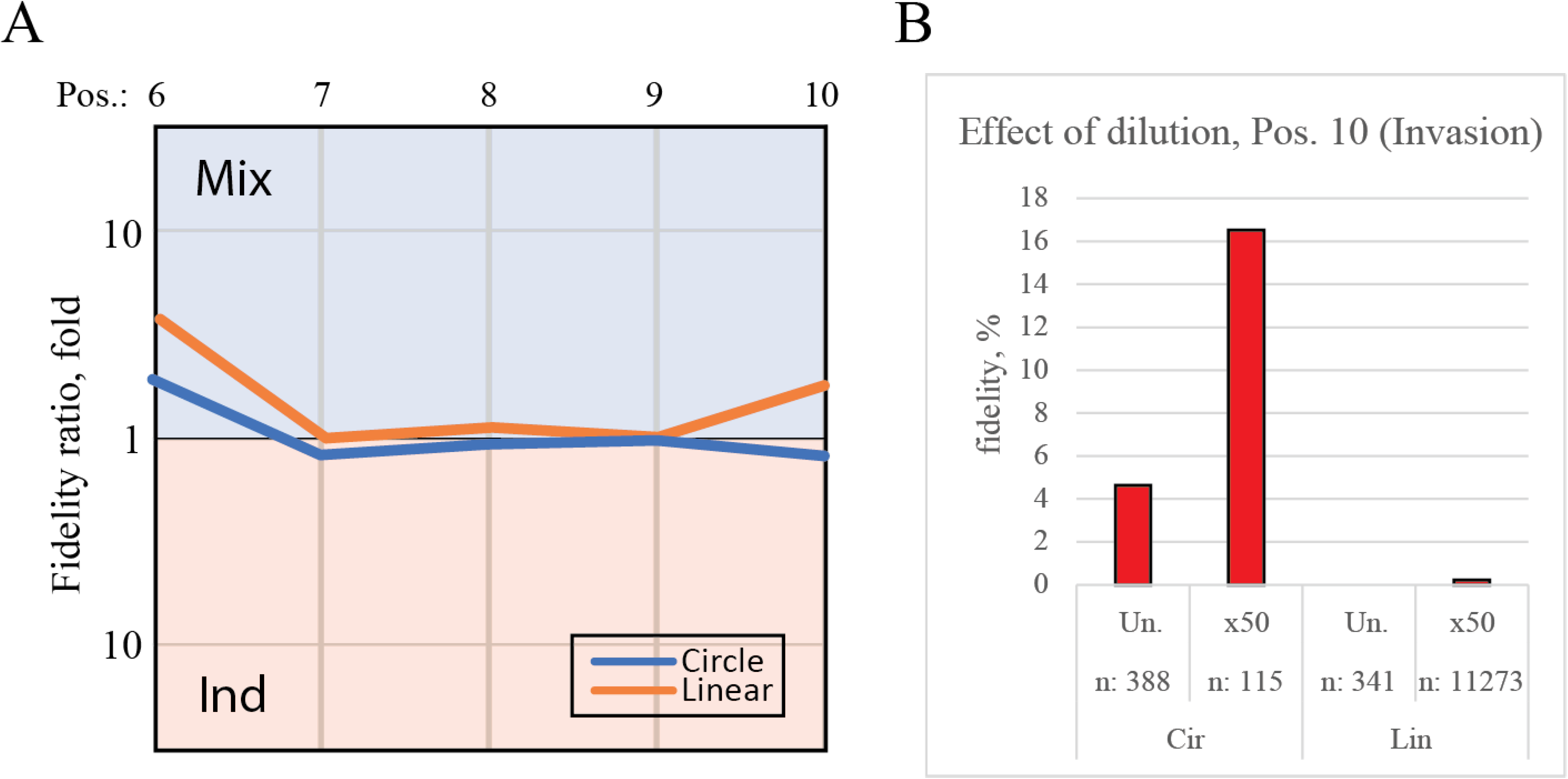
Controls for deep-sequencing data. A) Plot shows the fidelity ratio at the noted triplet positions between extension reactions where the templates were incubated either in one-pot (Mix) or in individual tubes (Ind). This is shown for the circular templates (blue lines) and linear templates (orange line). B) The effect of dilution with water leads to increased fidelity at the point of invasion (position 10). Bar chart show the fidelity for insertion of the expected triplet at position 10 (making invasion) calculated from deep-sequenced samples that were either not diluted (Un.) or diluted 50 fold (x50).

**Figure 5 - Figure supplement 1.**
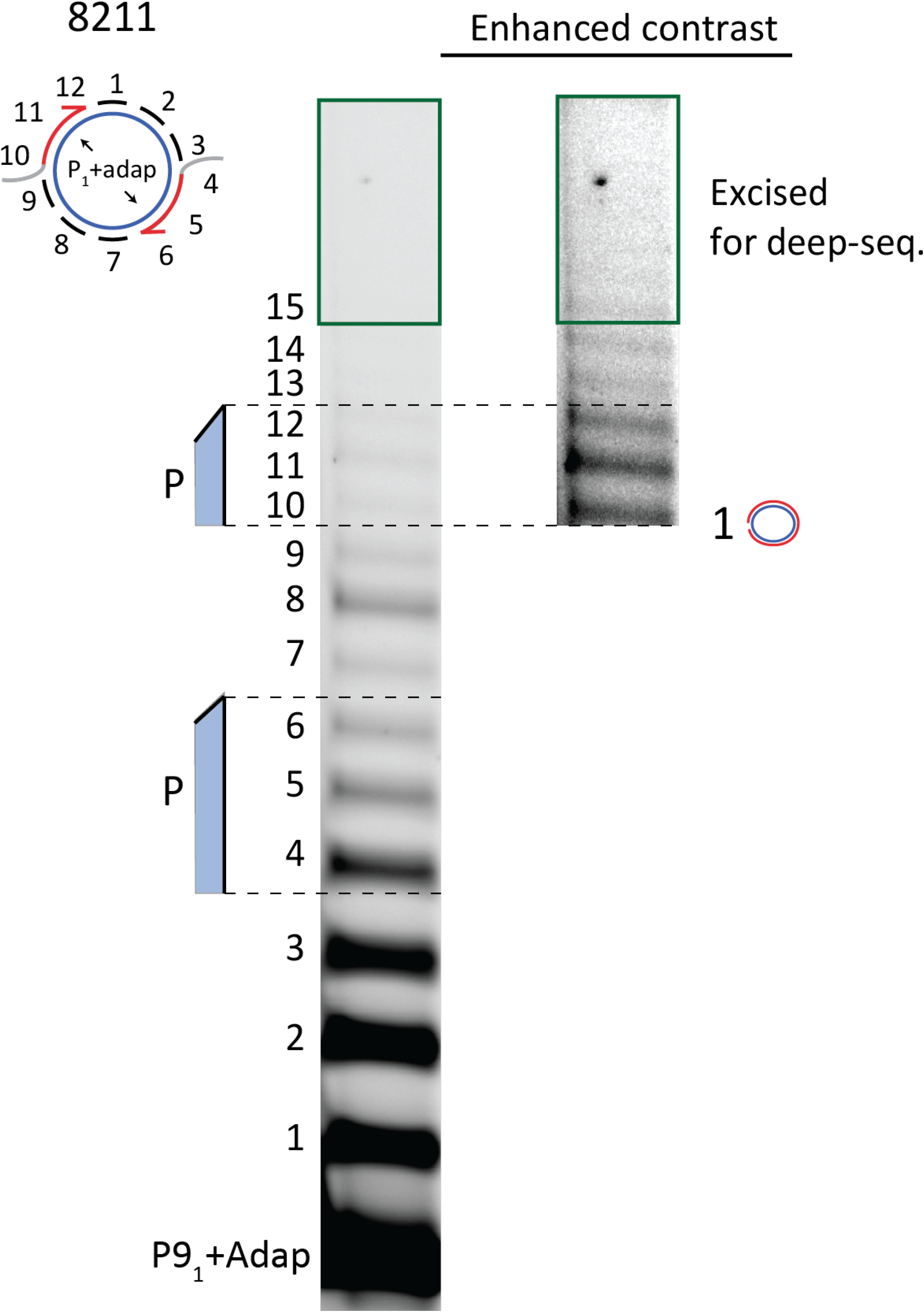
10% Urea PAGE separation of circular template extension reaction used for deep sequencing. Excised gel piece is marked with green.

**Figure 6 - Figure supplement 1.**
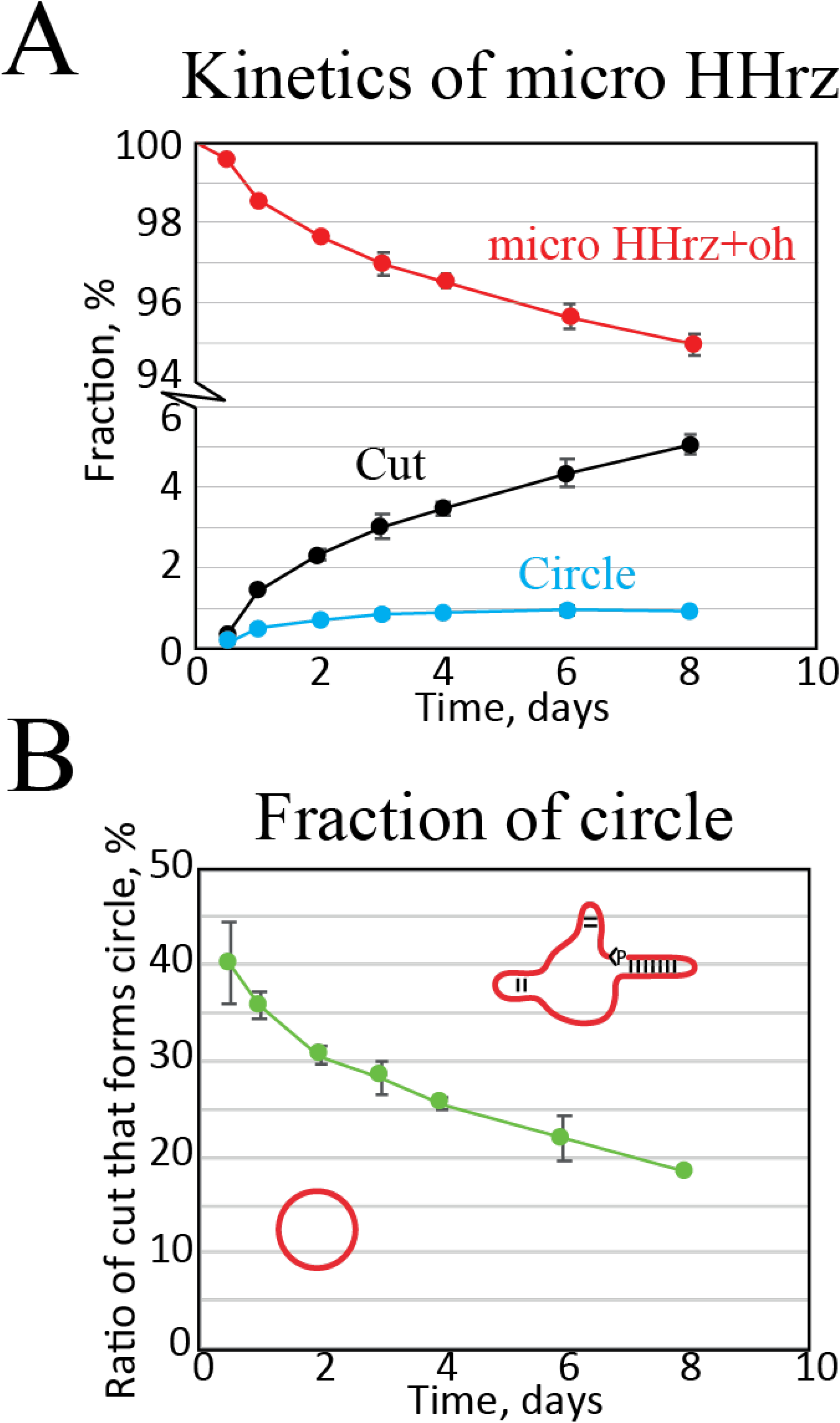
kinetic analysis of the micro HHrz. A) Quantification of band intensities as a function of time. Here we see that cut RNA accumulate while the amount of circular RNA seems to reach an equilibrium. B) Fraction of circular RNA relative to cut as a function of time. This plot shows that a very high amount of circle is formed at short time points slowly dropping in relation to non-circular cut RNA.

**Supplementary Movie 1.** Movie of the RCS simulation where dsRNA is 27 bp long. We observe fraying and annealing of 5’ and 3’ ends demonstrating the quick timescales of these transitions.

**Supplementary Movie 2.** Movie of the RCS simulation where dsRNA is 30 bp long. We observe again fraying and annealing of 5’ and 3’ ends demonstrating the quick timescales of these transitions.

